# CellPulse: A Foundation Model of Coordinated Gene Dynamics Simulating Viral Infectious Diseases

**DOI:** 10.64898/2026.04.22.720078

**Authors:** Dan Liu, Xiao-Xu Zhu, Libo Zhang, Deyu Xu, Jing Lou, Xiaobei Xiong, Yujie Ren, Yanjun Wu, Xi Zhou

**Author notes:** Corresponding authors: Libo Zhang, Yujie Ren, Yanjun Wu, Xi Zhou. The authors contribute equally (Dan Liu, Xiao-Xu Zhu, Libo Zhang).

## Abstract

Understanding how cells respond to perturbations like viral infections requires models capturing coordinated gene dynamics. However, current gene expression foundation models are predominantly reliant on single-cell data and static gene expression, limiting their applicability in real clinical scenarios. We present CellPulse, a direction-aware foundation model trained on the Virus Stimulated Atlas (VISTA), a newly curated atlas of over 23 million bulk RNA-sequencing differential expression profiles from viral infections. CellPulse models the direction and magnitude of gene expression changes via a structured representation of differential expression and a direction-aware attention mechanism, enabling the learning of coherent regulatory programs. It shows powerful diagnosing capability by accurately classifying 31 distinct virus types across diverse clinical and laboratory samples, solely from host transcriptional signatures. Crucially, without prior knowledge injection, CellPulse’s interpretability reveals virus-associated host factors that mediate infection. Using a selection of host factors for *in silico* drug screening yielded numerous compounds with confirmed efficacies in wet-lab assays, while cell-based and animal experiments further verified the causal relationship between host targets and viral infections. Overall, CellPulse represents a generalizable foundation model for deciphering coordinated gene dynamics from bulk transcriptomics, bridging host response modeling with clinical relevance and therapeutic discovery for infectious diseases and beyond.

## Introduction

Understanding how biological systems respond to external perturbations, such as viral infections, chemical exposures, or environmental stressors, remains a central challenge in systems biology and medical researches. Transcriptome-wide profiling, particularly through RNA sequencing (RNA-seq), has become an indispensable tool for quantifying cellular responses under such conditions^1^. Differential gene expression analysis allows researchers to identify genes whose expressions are altered between conditions^2–6^. However, these analyses often remain limited to static gene-level summaries or fold-change rankings. Critically, existing computational frameworks frequently disregard the contextual and directional nature of these changes, overlooking the coordinated regulatory patterns that underlie cellular adaptation and disease progression.

Recent efforts to model transcriptomic data through deep learning have made significant advances. Models such as Geneformer^7^, GeneCompass^8^, CellPLM^9^, Cell2Sentence^10^, scGPT^11^ and scBERT^12^ adopt language modeling strategies to embed genes and learn co-expression patterns across cellular contexts. These models have demonstrated impressive performance in transfer learning tasks and cross-condition prediction. Yet these approaches are primarily trained on raw expression matrices or gene co-occurrence signals in static contexts, limiting their capacity to generalize to dynamic perturbation scenarios, especially when the direction and intensity of regulation carry crucial biological meaning. Besides, these models are predominantly trained on single-cell RNA sequencing (scRNA-seq) data, which limits their usage in real clinical scenarios^13^. On the other hand, bulk RNA-seq provides much higher sequencing depth and lower dropout bias, can measure more subtle gene expression changes, and is more compatible with diverse laboratory and clinical samples (even including frozen or fixated ones), technically easier to perform, and much more cost-effective for ordinary medical examinations. However, despite of these advantages, comparing with scRNA-seq that can generate thousands of expression profiles in a single sample, bulk RNA-seq only generates one expression profile per sample, dramatically limiting its ability to produce large quantity of datasets for large model training^14,15^. Thus far, there is no model has systematically learned disease- or perturbation-relevant representations from bulk RNA-seq data that are more widely available and clinically relevant.

To address this limitation, we require a large-scale, diverse, and biologically relevant database of transcriptional responses to various viral infections for training a large foundation model. For this purpose, we constructed the VIrus STimulated Atlas (VISTA) by systematically processing 4,142 publicly available RNA-seq datasets from the Gene Expression Omnibus (GEO)^16,17^. Through exhaustive pairwise comparisons between experimental samples, we generated over 23 million differential expression (DE) profiles spanning 43,579 human genes. This process not only generates biologically meaningful gene DE profiles, but also highly expands the data quantity for further model training. To further specialize VISTA toward modeling virus-host dynamics, we manually curated 972 GEO series with viral infection metadata, including taxonomy from 52 viral types and critical experimental variables such as dose and time post-infection. This resulted in a unique corpus of 577,011 viral perturbation profiles, forming, to our knowledge, the largest labeled resource of its kind.

Based on the VISTA, we developed CellPulse, a direction-aware foundation model designed to decode transcriptional responses to external stimuli, with a specific focus on virus-host interactions. Unlike prior static embedding models, CellPulse is tailored to ingest gene-level differential expression profiles, capturing not only the relative strength of gene regulation through magnitude-based ranking, but also the direction of each gene’s regulation. Explicit modeling of regulatory direction is critical, as opposite changes of the same gene may correspond to fundamentally distinct biological states and pathogenic mechanisms. Treating differential expression as direction-agnostic signals risks conflating activation with repression, thereby obscuring meaningful regulatory contrasts. By preserving both regulatory strength and polarity, CellPulse captures the full semantics of transcriptional perturbations. Through a structured encoding of magnitude-ranked gene sequences together with directional regulatory signals and a direction-aware attention design, CellPulse learns coherent gene co-regulation programs across diverse perturbations. This enables the model to move beyond static expression patterns and capture the underlying organizational principles of coordinated transcriptional responses.

CellPulse was first pre-trained in a self-supervised manner^16^ using the full VISTA dataset, optimizing a masked gene prediction objective inspired by masked language modeling in BERT^17^. This stage enables the model to learn generalizable representations of gene co-regulation patterns across diverse perturbations without requiring labeled endpoints. In a subsequent fine-tuning stage, CellPulse was trained with virus identity labels to discriminate among different viral infections based on host differential expression profiles. Through this process, the model learns to capture virus-associated host transcriptional response patterns, which can be interpreted as virus-specific transcriptional signatures, i.e. Virus Signatures, embedded in the latent space. Leveraging these learned representations, CellPulse accurately identifies viral taxonomy using host gene expression responses alone, establishing a host response-based framework for viral infection diagnosis.

Beyond the classification/diagnosis performance, CellPulse provides a principled mechanism for model interpretability. By leveraging attention-based attribution methods^18^, the model enables direct identification of viral infection-associated host factors (VAHFs), whose transcriptional responses contribute most strongly to viral discrimination. These VAHFs include not only a large number of genes in well-established virus-relevant pathways but also numerous ones without previously reported link with viral infections, which highlight host pathways and regulatory components that are closely associated with viral perturbations and therefore represent candidate host factors for therapeutic exploration. By integrating the identified VAHFs with compound-target knowledge bases, we further enable *in silico* prioritization of antiviral compounds. The resulting candidates were experimentally evaluated in selected viral systems using both cellular and *in vivo* models, demonstrating the translational potential of the framework.

In summary, VISTA provides a large-scale and disease-relevant AI-ready database of host gene alterations responding to diverse viral infections, and CellPulse offers a powerful and generalizable framework for modeling host transcriptional responses under perturbation (Fig. S1). By combining large-scale differential expression data with an interpretable and biologically grounded architecture, CellPulse facilitates the systematic analysis of perturbation-induced host regulatory programs and advances our ability to decode the complex molecular logic of host responses to infections and probably many other diseases, providing a foundation for both mechanistic discovery and therapeutic prioritization.

## Results

### Constructing a large-scale perturbation atlas of host transcriptomic responses to viral infection

To support large-scale modeling of host transcriptional responses to viral infection, we constructed an AI-ready resource termed VISTA (Fig. 1a). We systematically retrieved 14,021 virus-related bulk RNA-seq studies (GEO Series, GSEs) from the NCBI GEO database using "virus infection" as the keyword. After applying stringent quality control criteria to exclude non-expression profiling assays (e.g., methylation data) and studies profiling fewer than 5,000 genes, we retained 4,142 high-quality virus-related GSEs comprising 158,370 individual samples (GSMs). These studies span a wide range of biological contexts, including clinically derived specimens such as whole bloods, peripheral blood mononuclear cells (PBMCs), isolated blood cells, nasopharyngeal swabs (NP swabs), and tissue biopsies, as well as laboratory-generated infection samples from multiple cell lines and mouse models.

**Figure 1.**
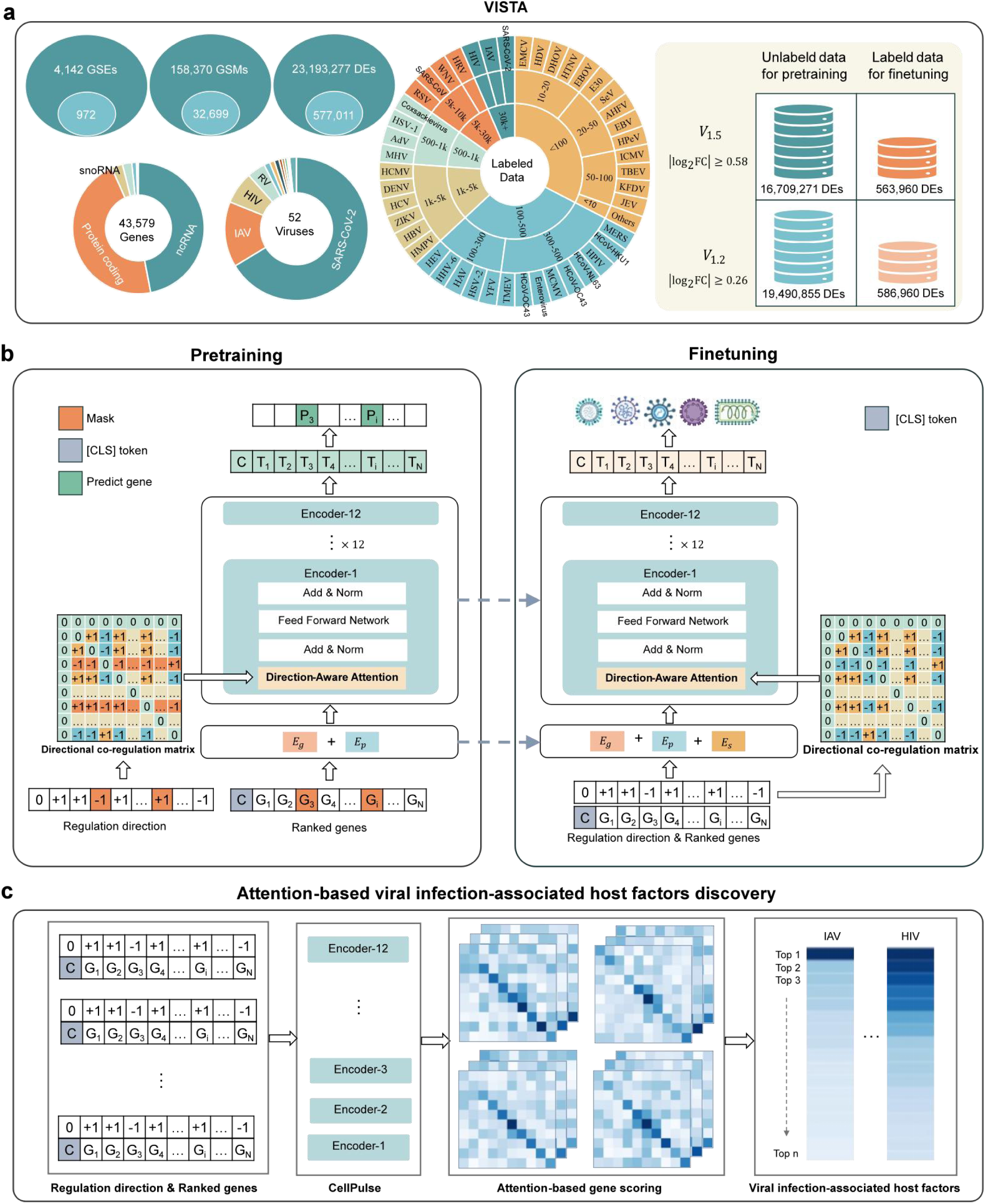
VISTA dataset and CellPulse architecture. **a,** Overview of the VISTA dataset. VISTA integrates 4,142 GSEs, comprising 158,370 samples and over 23 million differential expression (DE) profiles across 43,579 human genes. A curated subset of 972 GSEs, including 32,699 samples and 577,011 DE profiles, is annotated across 52 viral types. Two AI-ready versions, _1.5_ and _1.2_ , are provided. **b,** Schematic of CellPulse pretraining and fine-tuning. Each DE profile is encoded as a ranked gene sequence with associated regulation directions. CellPulse employs 12 stacked Transformer encoder layers in which the standard self-attention module is replaced with a direction-aware attention mechanism that highlights co-regulated genes. During fine-tuning, CellPulse transfers its pre-trained regulatory representations for identification of virus-specific host transcriptional signatures. **c,** Discovery of viral infection-associated host factors (VAHFs). Attention weights learned by CellPulse are analysed to identify key host factors associated with distinct viral infection responses.

To ensure comparability across samples and species, raw expression data were harmonized into normalized expression matrices with all genes standardized to Entrez IDs. Mouse transcriptomes were further aligned to human gene space through ortholog mapping, yielding unified expression matrices suitable for integrative and cross-species analyses.

A central challenge in aggregating public infection transcriptomes is the heterogeneity of experimental designs and the frequent absence or inconsistency of explicit control annotations. To maximize usable regulatory signal while minimizing dependence on curated metadata, we performed exhaustive pairwise differential expression analyses within each study. Specifically, for any two samples measured within the same GSE, we computed gene-level log_2_ fold changes, treating each sample alternately as the reference. This strategy generates differential expression (DE) profiles without assuming predefined case–control groupings and directly extracts regulatory shifts from expression contrasts. Each DE profile thus encodes a biologically coupled transcriptional response. This design rests on a simple but powerful biological principle: transcriptional regulation is inherently coordinated, and any differential expression profile, regardless of how it is constructed, reflects structured, coupled gene responses rather than isolated gene changes.

Using this strategy, we generated 23,193,277 DE profiles, spanning 43,579 human genes, including protein-coding genes (46.318%), non-coding RNAs (ncRNAs, 46.993%), and other gene categories. In addition, to support virus-specific learning, we manually annotated 972 virus-related GSE entries with viral taxonomy (52 virus types) and key infection parameters (dose and time). This curation resulted in 577,011 virus-associated DE profiles, making VISTA the most comprehensive resource of host transcriptomic responses to viral infection constructed to date.

From the full set, we derived two AI-ready datasets that share identical construction logic but differ in fold-change stringency. Applying a 1.2-fold threshold resulted in the dataset *V*_1.2_ , comprising 19,490,855 unlabeled DE profiles and 586,960 virus-associated DE profiles, whereas a more stringent 1.5-fold threshold produced *V*_1.5_ , containing 16,709,271 unlabeled DE profiles and 563,960 virus-associated DE profiles. These two datasets form parallel yet biologically distinct foundations for evaluating the robustness of CellPulse under different perturbation intensities. The resulting VISTA dataset thus serves both as a large-scale pretraining resource for representation learning and a unified benchmark for downstream viral perturbation modeling, enabling a comprehensive assessment of how molecular response patterns encode pathogen-specific regulatory programs.

### Learning gene co-regulation patterns

To move beyond static expression modeling, which treats gene expression as isolated measurements and struggles to capture coordinated regulatory behavior, we developed CellPulse (Fig. 1b), a Transformer-based architecture tailored to learn gene co-regulation patterns from perturbation transcriptomes. The design builds on a key biological observation: differential expression (DE) profiles are inherently determined by coordinated gene responses, because genes rarely shift individually but instead change in structured, interdependent patterns. Modeling these relative shifts, rather than absolute abundance, therefore provides a more direct window into the regulatory logic governing cellular responses.

Motivated by this principle, CellPulse operates on differential expression signals derived from RNA-seq contrasts, transforming each DE profile into a structured representation composed of two complementary components: (1) a ranked gene list sorted by absolute log_2_ fold-change (|log_2_FC|), prioritizing genes with the largest expression deviations; and (2) a direction vector indicating up- or down-regulation for each gene. This design treats each DE profile as a gene sequence, where genes act as "tokens" arranged by the magnitude of their transcriptional deviation. Unlike natural language tokens^19^, each gene token additionally carries an explicit regulation direction (up- or down-regulation), allowing the model to jointly encode magnitude, polarity, and contextual co-regulation, aligning naturally with how biological systems orchestrate perturbation responses and conferring advantages over models that rely solely on static expression levels. It is noteworthy that the name of this model, CellPulse, precisely reflects its nature to model the gene co-regulation dynamics in cells (i.e. Genes’ Pulse in Cells).

We adopted a self-supervised pre-training strategy based on contextual masking to learn gene co-regulation patterns directly from large-scale perturbation data without relying on explicit labels. During pre-training, 15% of gene tokens were randomly masked, and CellPulse was trained to recover their identities from the surrounding genes in the same differential expression profile. By predicting masked genes within ranked perturbation contexts, the model is encouraged to capture recurrent gene co-regulation patterns that characterize coordinated transcriptional responses. A central challenge of pairwise DE profiles is that regulation direction ("up" or "down") is inherently relative and lacks a global reference across datasets. To address this, we introduce a directional co-regulation matrix *S* ∈ { − 1,0, + 1}*^N^*^×*N*^, encoding concordant ( +1 ) or discordant ( −1 ) regulatory trends between gene pairs while zeroing self-relations. This matrix was integrated into the self-attention mechanism to prioritize interactions between genes sharing coherent regulatory behavior while attenuating attention across opposing trends. By embedding directional co-regulation as an inductive bias within attention, CellPulse is able to learn higher-order coordination patterns that conventional expression-only Transformers cannot capture^7–12^.

Through this integrated design, CellPulse acquires direction-aware, context-sensitive gene embeddings that capture stable co-regulation patterns across a wide range of host-pathogen interactions. These embeddings provide a structured representation of how genes jointly respond to perturbations, forming the foundation for downstream tasks including viral identity inference, host-dependency gene discovery, and mechanistic interpretation of infection-induced transcriptomic shifts.

### Diagnosing viral infection identities by CellPulse

Diagnosing viral infection types in real-world clinical samples is very challenging. To evaluate the biological relevance and diagnostic power of CellPulse representations in infectious contexts, we applied the model to predict virus types in diverse clinical and laboratory samples based on host transcriptomic perturbations.

Because CellPulse is pretrained to capture coordinated up/down-regulation patterns across diverse perturbations, this task naturally tests its transferability: if the learned representations truly encode generalizable co-regulation principles, they should support accurate discrimination of viral infections solely from corresponding host responses, regardless of experimental conditions or virus types. For comparison, we included a BERT baseline that employs standard Transformer encoders operating on the same ranked gene sequences but without explicit modeling of regulation direction or directional co-regulation patterns. This baseline serves to assess the contribution of CellPulse’s direction-aware design beyond generic sequence modeling capacity. To ensure a fair evaluation and avoid bias introduced by threshold choice, we selected the intersection of 1.2-fold dataset (*V*_1.2_) and 1.5-fold dataset (*V*_1.5_) and retained 31 virus types with at least 100 DE profiles each, resulting 555,151 examples. Besides, 130,191 uninfected examples (generated by mock-mock sample pairs) were included. These were then divided into training, validation, and test sets at an 8:1:1 ratio (548,270; 68,536; and 68,536 examples), providing a rigorous benchmark for assessing how effectively CellPulse transfers its self-supervised knowledge to real-world viral infection prediction.

Fine-tuning the pre-trained CellPulse model on the VISTA benchmarks produced consistently strong classification performance on both *V*_1.5_ and *V*_1.2_ . Because viral classes are highly imbalanced in quantities, model performance was evaluated using macro-F1. Macro-F1 is the arithmetic mean of per-class F1 scores (the harmonic mean of precision and recall computed for a single class) and thus reflects the model’s ability to maintain balanced precision-recall performance across all classes. On *V*_1.5_, CellPulse reached a macro-F1 of 0.854, compared to 0.840 for the BERT baseline. The performance gap widened on the larger *V*_1.2_ dataset, where CellPulse achieved 0.908, exceeding BERT’s 0.882 (Fig. 2a). The corresponding normalized confusion matrices further illustrate that these gains arise from reduced cross-class confusion and more balanced predictions across viral categories (Fig. S2 and S3). These consistent improvements indicate that modeling up/down regulatory direction substantially enhances the ability to decode virus-specific transcriptomic signatures. Per-class F1-scores further highlight these advantages (Fig. 2b). Well-characterized viruses, including Severe Acute Respiratory Syndrome coronavirus-2 (SARS-CoV-2), Influenza A virus (IAV), rhinovirus, West Nile virus (WNV), human cytomegalovirus (HCMV), Dengue virus (DENV), Hepatitis B virus (HBV), mouse hepatitis virus (MHV), mouse cytomegalovirus (MCMV), HCoV-HKU1, MERS-CoV, and herpes simplex virus-1 (HSV-1) and -2 (HSV-2), achieved high F1-scores (> 0.90) under both models and thresholds, reflecting strong, distinctive transcriptomic signatures. In contrast, several challenging virus classes showed marked improvements only when using CellPulse. For example, under the more stringent *V*_1.5_ condition, BERT obtained F1-scores of 0.655 for adenovirus and 0.716 for enterovirus, whereas CellPulse increased these values to 0.679 and 0.730, respectively. These gains were even more pronounced under the *V*_1.2_ threshold, such as adenovirus (0.875 vs. 0.921) and enterovirus (0.847 vs. 0.870). Importantly, CellPulse also enhanced prediction for the uninfected class, which is quite challenging due to inherent heterogeneity in baseline gene expressions of various samples. On *V*_1.5_ , BERT achieved an F1 of 0.720, whereas CellPulse improved this to 0.842. On *V*_1.2_, the improvement persisted (0.757 vs. 0.907), demonstrating that direction-aware modeling not only captures virus-specific perturbations but also distinguishes normal, uninfected profiles with high fidelity. These results show that CellPulse reliably prioritizes biologically meaningful transcriptional signals, enhancing classification across both strong and weak viral responses, and improves identification of uninfected host states, reflecting its capacity to model coordinated regulatory direction in complex transcriptomic landscapes.

**Figure 2.**
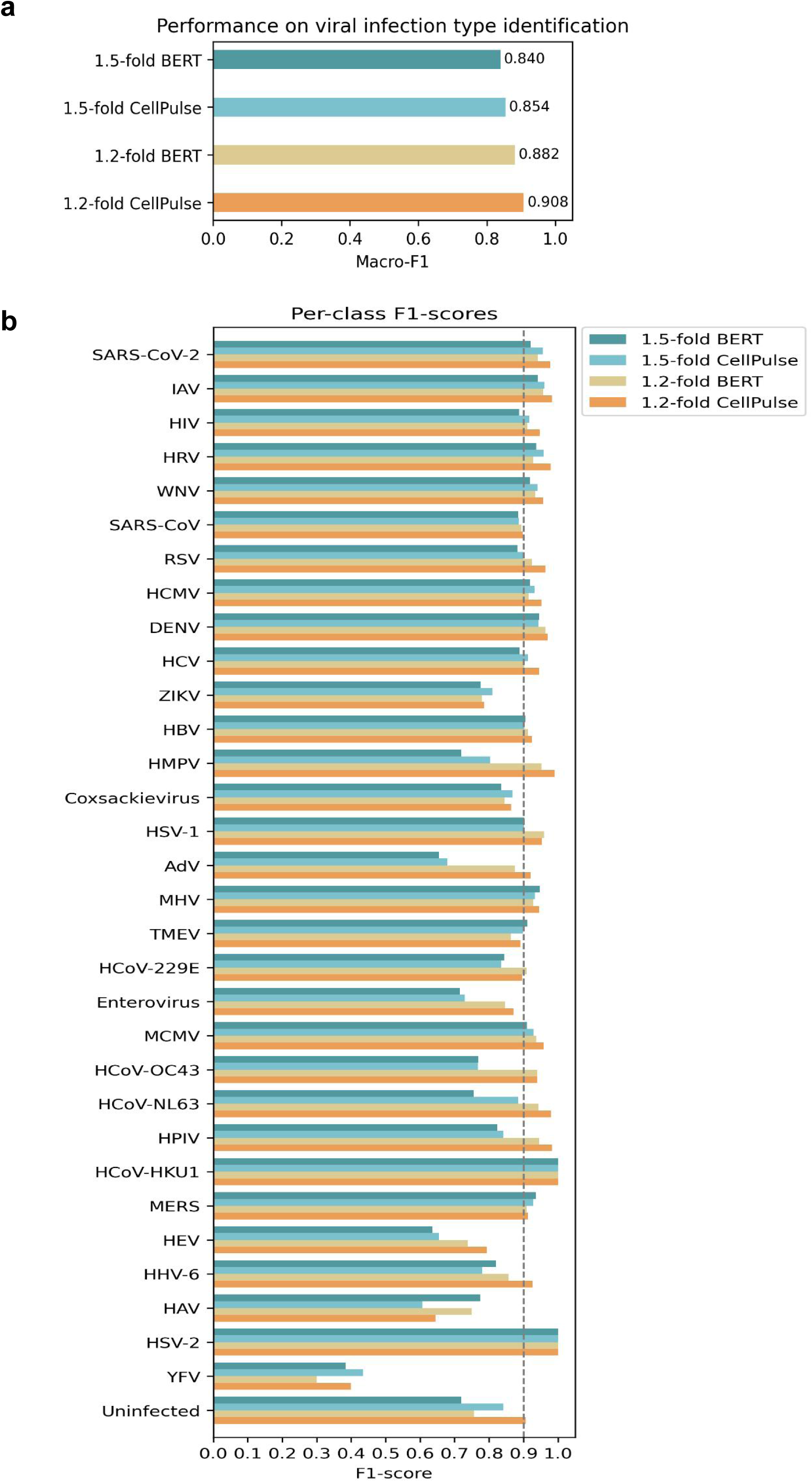
Comparison of macro-F1 and per-class F1 scores between the BERT baseline and CellPulse under 1.5-fold and 1.2-fold thresholds. **a**, Performance on viral infection type classification. Macro-F1 scores, reflecting balanced precision–recall performance across all classes, are shown for the BERT baseline and CellPulse under the indicated thresholds. **b,** Per-class F1 scores for each virus under the indicated thresholds, comparing the BERT baseline and CellPulse.

Interestingly, the comparison between the two thresholds revealed an important biological insight. Although the 1.2-fold dataset ( *V*_1.2_ ) includes a larger number of genes with more subtle expression change and therefore introduces more background variation and potential noise, CellPulse demonstrated superior predictive performance under this condition. This suggests that the model effectively disentangles biologically informative co-regulation signals from background fluctuations, selectively emphasizing genes that carry virus-specific transcriptional signatures, i.e. Virus Signatures. The improved performance under a more permissive threshold indicates that viral infection triggers distributed transcriptional programs beyond the most strongly perturbed genes, and that the model is capable of integrating these subtle yet coordinated changes to infer virus signature and viral identity.

Collectively, these findings demonstrate that CellPulse not only captures the dominant expression shifts associated with infection but also leverages fine-grained regulatory patterns embedded within broader transcriptional responses. Direction-aware modeling further enhances classification of challenging virus classes, underscoring the importance of encoding transcriptional directionality for decoding complex host-pathogen interactions.

### Distinct influenza subtypes can be distinguished by CellPulse

Beyond discriminating the identities of distinct virus types, we further assessed whether CellPulse can resolve finer-grained viral subtypes that share substantial biological similarity. IAV represents a particularly challenging test case, as different subtypes often induce overlapping host transcriptional responses while showing some difference in pathogenicity, host or cell/tissue tropism, and immune evasion mechanisms. To this end, we extracted all IAV-related DE profiles from VISTA and manually curated subtype annotations by tracing the original experimental metadata. After excluding subtypes with fewer than 100 DE profiles to ensure statistical robustness, five IAV subtypes were retained: H1N1, H3N2, H5N1, H7N9, and H7N7. The resulting dataset exhibits substantial class imbalance (Fig. 3a) and spans both seasonal and avian IAV strains, posing a stringent evaluation scenario.

**Figure 3.**
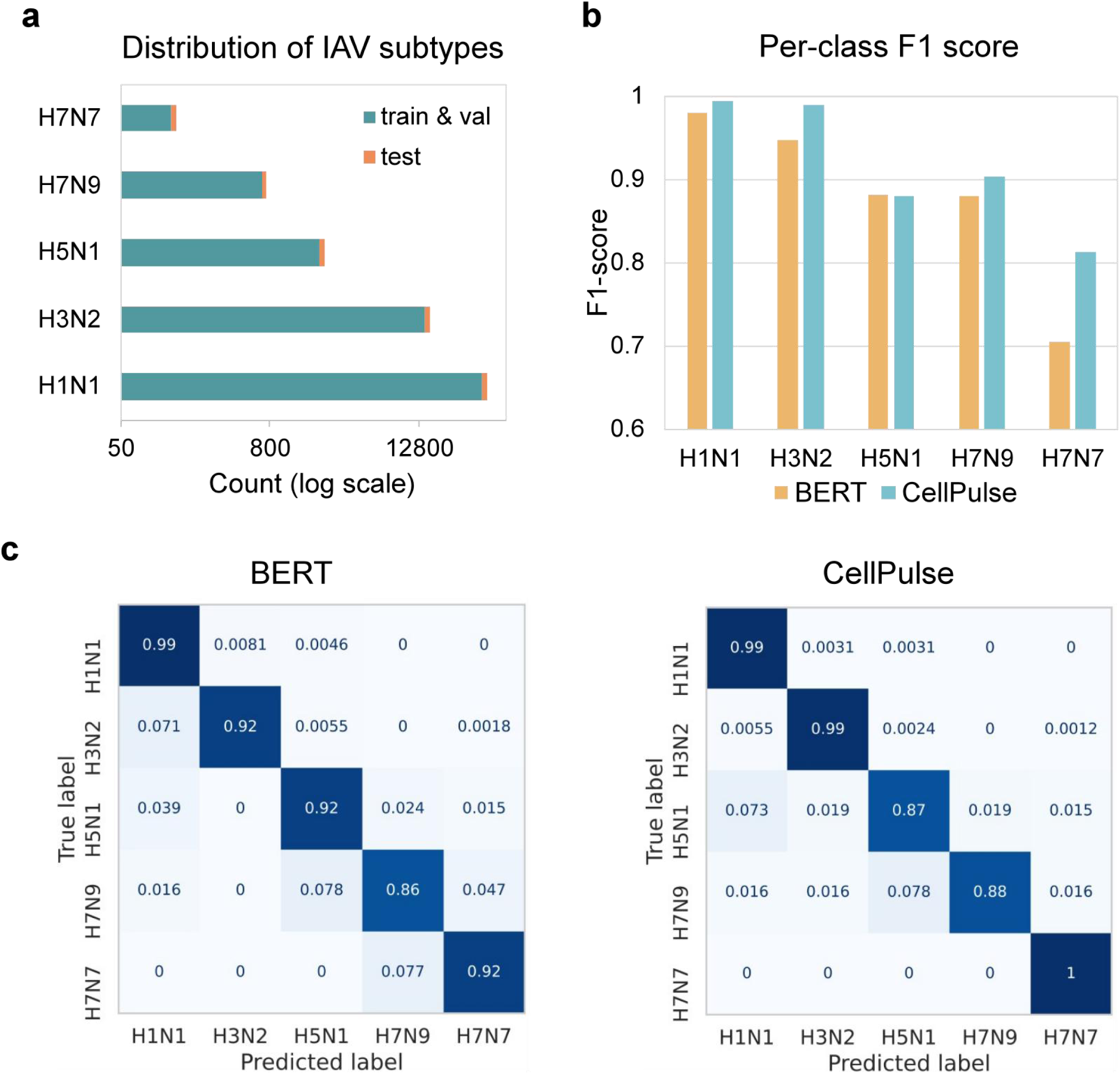
Fine-grained IAV subtype classification using CellPulse. **a**, Distribution of IAV subtypes. **b,** Per-class F1 scores on the held-out test set for BERT and CellPulse. **c,** Confusion matrices of BERT and CellPulse on the test set, normalized by true subtype counts (row-wise). Left: BERT; Right: CellPulse.

Profiles were randomly partitioned into training, validation, and test sets using an 8:1:1 ratio. Starting from the pre-trained CellPulse encoder, we fine-tuned the model for multi-class IAV subtype classification. Model performance was evaluated on the held-out test set using macro-F1 score, which equally weights all subtypes and is robust to class imbalance. The same evaluation protocol was applied to the BERT baseline described above.

CellPulse achieved a macro-F1 score of 0.9159, exceeding the BERT baseline (0.8792), reflecting an overall improvement in influenza subtype discrimination under severe class imbalance (Fig. 3a). Per-class F1-score comparisons indicate that the performance gain is not uniformly distributed across subtypes (Fig. 3b). For highly represented or transcriptionally distinctive subtypes such as H5N1, CellPulse and BERT exhibit nearly identical F1 scores, suggesting that both models can reliably capture strong and characteristic host response patterns. In contrast, CellPulse shows clear advantages for low-frequency subtypes, including H7N9 and H7N7, where classification is more susceptible to noise and cross-subtype confusion. This pattern is further supported by confusion matrix analysis (Fig. 3c). While BERT yields a clean diagonal for dominant or strongly separable subtypes, it exhibits more diffuse off-diagonal predictions for minority classes. CellPulse produces a more balanced confusion structure, with reduced misassignment of rare subtypes into dominant categories, indicating improved subtype-specific separation rather than overconfident predictions driven by class prevalence. Thus, such an advantage makes CellPulse an ideal model to study newly emerging influenza subtypes.

These results demonstrate that CellPulse captures subtle yet systematic differences in host regulatory responses induced by closely related viral subtypes. This capability extends CellPulse’s applicability from diagnosing virus types to fine-grained viral subtype resolution, underscoring its potential for detailed pathogen characterization and surveillance based on host transcriptomic responses.

### CellPulse shows superior diagnosing power in different types of viral infection samples

As aforementioned, VISTA includes the bulk RNA-seq datasets derived from a variety of clinical and laboratory samples. Since CellPulse achieved high predictive accuracy for most types of viruses, it is intriguing to examine how the model performs in different types of samples. We used the CellPulse model trained on _1.2_ that shows the best performance in predicting virus types. As shown in Fig. 4a, CellPulse achieved high predictive accuracies for both clinical and non-clinical laboratory samples, as an accuracy of 96.72% for the clinical datasets and 86.70% for the laboratory ones. Within the clinical samples, the predictive accuracy reached accuracy of 98.80% for blood samples (i.e. whole blood), which are the most common clinical samples in routine physical examinations and standard medical testings and comprised of ∼60% clinical samples in VISTA, and 93.63% for non-blood samples, respectively (Fig. 4b). These results demonstrate robust generalizability across data sources.

**Figure 4.**
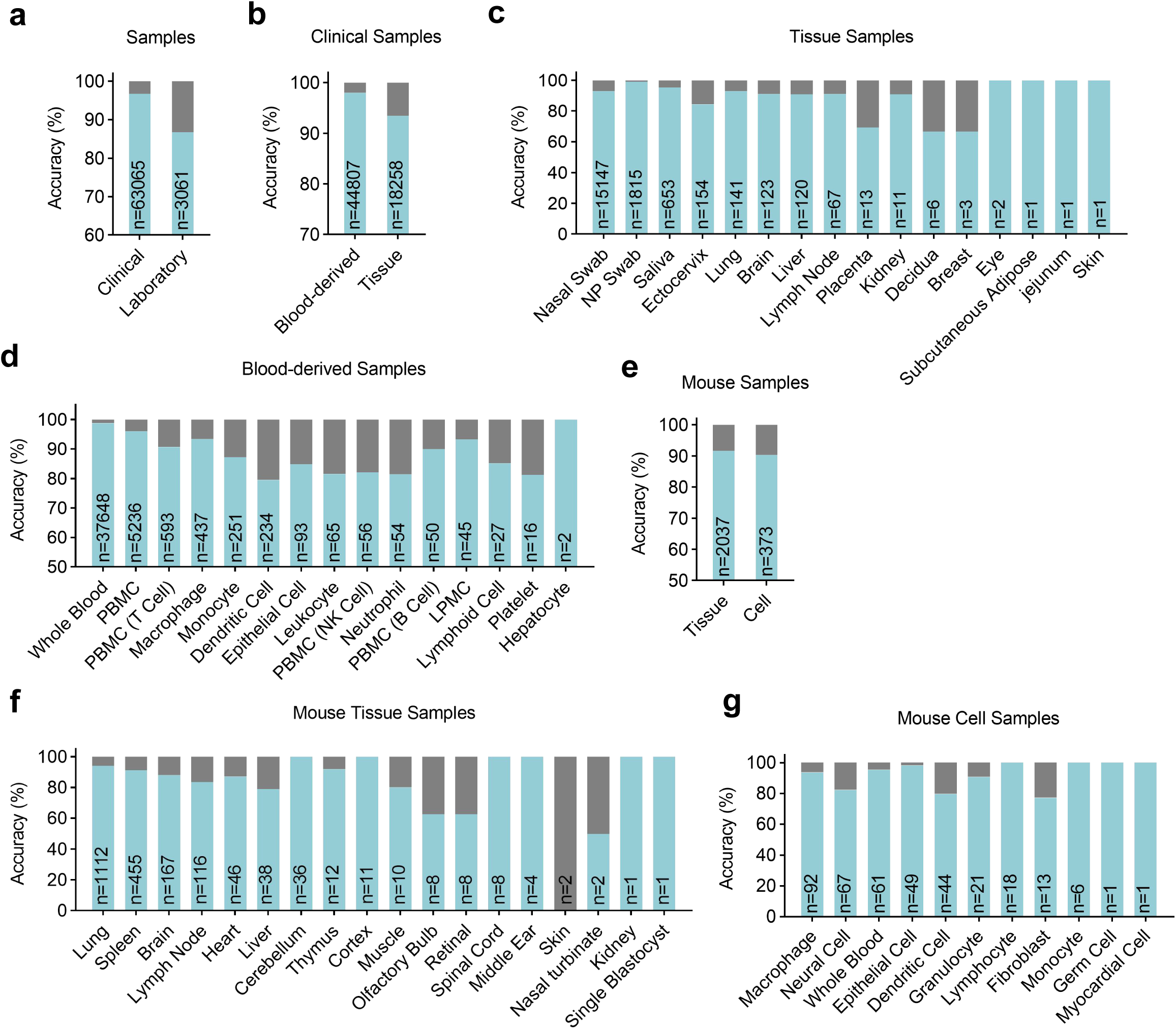
CellPulse shows superior diagnosing power in different types of viral infection samples. **a**, Diagnostic accuracy of CellPulse in human-derived viral infection samples from clinical (*n* = 63,065) and laboratory (*n* = 3,061) settings. **b,** Diagnostic accuracy of CellPulse in blood-versus tissue-derived clinical viral infection samples (*n* = 44,807 and 18,258, respectively). **c,** Diagnostic accuracy of CellPulse across human tissue types in clinical viral infection samples (*n* values shown above bars). **d,** Diagnostic accuracy of CellPulse across human blood-derived cell types in clinical viral infection samples (*n* values shown above bars). **e,** Diagnostic accuracy of CellPulse in murine-derived viral infection samples from tissue (*n* = 2,037) and cells (*n* = 373). **f,** Diagnostic accuracy of CellPulse across murine tissue types in viral infection samples (*n* values shown above bars). **g,** Diagnostic accuracy of CellPulse across murine cell types in viral infection samples (*n* values shown above bars).

To further delineate the CellPulse’s performance characteristics in different types of clinical samples, non-blood clinical samples were stratified by tissue origins including brain, breast, decidua, ectocervix, eye, kidney, liver, lung, lymph node, placenta, jejunum, subcutaneous adipose, nasal or nasopharyngeal swab (np swab), skin and saliva (Fig. 4c). Strikingly, the model showed strong class-wise predictive power, as accuracy exceeding 90% are achieved for nearly all the tissue types, with a substantial proportion achieving perfect 100% accuracy (Supplementary Table 1). Notably, the decidua and breast subgroups contain only 6 and 3 samples, respectively, as such small sample sizes are inherently vulnerable to accuracy fluctuations caused by individual misclassifications. Nevertheless, CellPulse still achieved a quite favorable predictive rate even in these small-sample cohorts, underscoring its robustness in diagnostics.

For clinical blood-derived samples, they were categorized by cell types, including, PBMCs, B cells, natural killer (NK) cells, T cells, dendritic cells, macrophages, leukocyte, monocyte, neutrophils, lymphoid cells, platelet, as well as hepatocyte, epithelial cells, Lamina Propria Mononuclear Cells (LPMCs) and whole blood sample (Fig. 4d). Except dendritic cells (accuracy of 79.5%), the predictive accuracy exceeded 80% for all the other cell types (Supplementary Table 2).

Furthermore, since VISTA contains some datasets from murine sources, we extended the evaluation to murine samples. Interestingly, these cross-species data achieved high predictive accuracy of 91.6% in total murine tissue samples and 90.35% in total murine cell samples (Fig. 4e), while most sub-categories, except ones with extremely small sample size, showed high accuracy, highlighting the cross-species applicability of CellPulse in translational and preclinical researches using mouse and probably other animal models (Fig.4f-g, Supplementary Table 3).

### Attribution and discovery of viral infection-associated host factors

To evaluate whether CellPulse captures biologically meaningful host responses and enables the systematic identification of VAHFs, we conducted a large-scale attribution analysis across different viral classes (Fig. 1c). Here, VAHFs are defined as host genes whose transcriptional responses are consistently and strongly associated with specific viral infections, reflecting host regulatory programs engaged during viral perturbation.

For each viral class, up to 10,000 DE profiles were randomly selected (using all available DE profiles if fewer than 10,000), resulting in a total of 76,038 DE profiles analysed. These profiles were processed by the fine-tuned CellPulse model to derive virus-specific transcriptional signatures, referred to as Virus Signatures.

To identify VAHFs from these signatures, we employed an attention weight-based attribution strategy that quantifies the contribution of individual genes to the model’s virus-specific predictions. Specifically, class-level gene importance scores were computed based on how strongly CellPulse attends to each gene when forming Virus Signatures for a given viral class. Genes with higher importance scores therefore represent host factors that the model relies on most heavily to discriminate a particular viral infection from others, providing a natural and interpretable measure of virus-associated gene relevance.

A key benchmark for evaluating AI-driven biological discovery is whether task-trained models can recover known biological determinants directly from data, without explicit supervision at the gene or pathway level. Accordingly, for each viral class, we ranked genes by their importance scores and selected the top 50 VAHFs (Supplementary Table 4). The top 50 VAHFs were further analysed by Gene Ontology (GO), a conventional bioinformatic method^20,21^. Strikingly, these CellPulse-identified VAHFs are enriched in GO categories related with viral infections, such as defense response to virus, humoral immune response, response to virus, response to chemokine, etc (Fig. 5), and include some well-established infection-related host factors in such as IFN-α2, IFN-α4, IFN-β1, ISG15, ISG20, USP18, IFIT1, IFIT2, IFIT3, IFITM3, MX1, MX2, IL-1β, IL-6, CXCL1, CXCL2, CXCL3, CXCL10, CXCL11, CCL2, CCL7, CCL8, TNF, TNFAIP6, etc across multiple virus types. We further characterized the functional properties of the CellPulse-identified VAHFs in the context of each individual virus. GO enrichment analysis was performed on the top 50 VAHFs for each virus (Fig. S4). The results revealed that, across a diverse panel of viruses, including Coxsackievirus, Enterovirus, HCMV, HHV-6, HIV, MCMV, Rhinovirus, SARS-CoV, SARS-CoV-2, TMEV, and WNV, the top 50 VAHFs were generally enriched in biological processes associated with viral infection, although distinct virus type showed some difference. These findings demonstrate the superior performance of CellPulse in rediscovering key host factors in fundamental pathways responsive to viral infections. Beyond recovering well-established host factors, CellPulse also prioritized a subset of genes not previously annotated as viral dependencies, marking them as high-confidence candidates for further experimental investigation.

**Figure 5.**
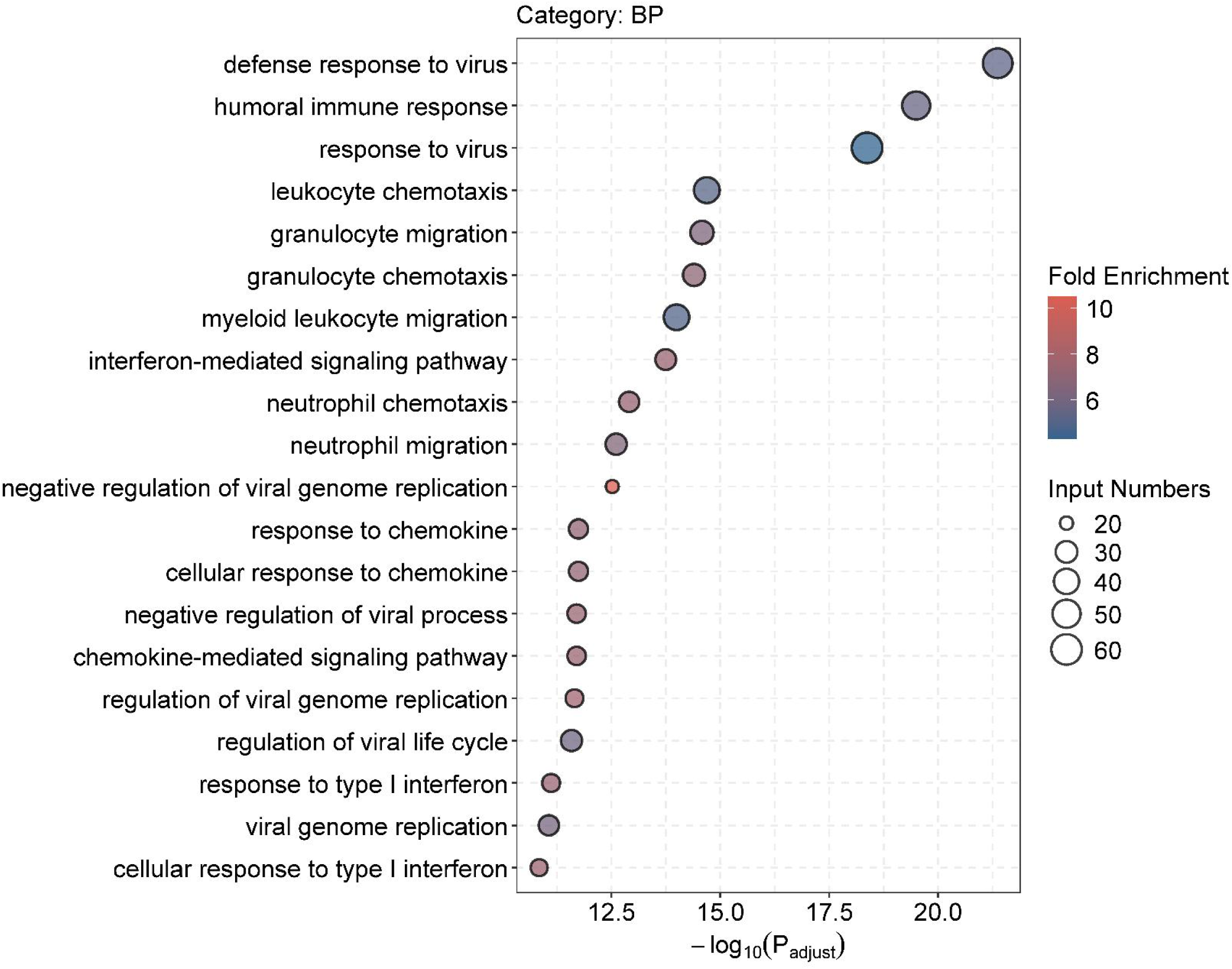
CellPulse-identified VAHFs are enriched in virus-relevant pathways. Bubble plot of the top 20 significantly enriched Gene Ontology (GO) on Biological Process (BP) terms associated with the gene set corresponding to the top 50 VAHFs across multiple viruses. GO enrichment analysis was conducted using a false discovery rate (FDR) threshold of < 0.05 for statistical significance. Bubble size represents the number of genes annotated to each term, and color intensity indicates the fold enrichment.

These results demonstrate that CellPulse not only diagnoses viral identities in infected samples with high accuracy but also exposes the underlying gene regulation circuitry orchestrating host-virus interactions. The attribution-based discoveries provide mechanistic interpretability for model decisions and offer a scalable path toward identifying conserved and virus-specific host targets with therapeutic potentials.

### Drug screening and validation based on VAHFs as potential targets

To translate VAHFs into actionable therapeutic targets, we mapped top 30 VAHFs for five representative viruses, including HSV-1 as a DNA virus, IAV as a negative-sense RNA virus, and ZIKV, DENV and MHV as positive-sense RNA viruses, to molecular targets archived in the ChEMBL database^22^. This was accomplished by first converting VAHFs to UniProt IDs via the UniProt’s ID mapping tool, followed by matching these UniProt IDs to their corresponding entries in ChEMBL^22^. This yielded a curated VAHF-target set that served as the basis for drug retrieval. Because these CellPulse-identified VAHFs include a number of well-established virus-related factors and even proven drug targets, we chose to exploit the potentials of the VAHFs without previous annotation of viral relevance as drug targets for viral diseases, which will provide novel insights into viral pathogenesis and serve as a good test for CellPulse’s performance.

Using these mapped targets, we identified candidate compounds by extracting all drugs in ChEMBL with at least one VAHF-derived target. Candidate compounds were subsequently ranked within each viral infection class based on two quantitative criteria: the number of VAHF targets modulated by each compound, and the corresponding activity values documented in ChEMBL database. Compounds targeting a greater number of VAHFs and exhibiting higher potency, i.e. lower half-maximal inhibitory concentration (IC_50_), were given higher priority. This ranking provides an unbiased, data-driven way to identifying drugs with the highest potential to modulate VAHF-associated host pathways. The analysis surfaced several compounds without prior links to the corresponding viral infections.

To further refine this compound library to ensure higher hit odds of candidate compounds, compounds with an IC_50_ exceeding 5000 nM, which was listed in ChEMBL, were excluded. Besides, for candidate compounds with the same target in ChEMBL, we sorted them in ascending order of IC_50_ values and selected the top 5 commercially available ones. If the number of qualified candidate compounds corresponding to a specific target was less than 5, all the eligible compounds were included in the candidate pool without further filtering. For each virus, candidate compounds were examined for their effects on viral replication in cells. In brief, HeLa cells, HEK293 cells and primary mouse lung fibroblasts (MLFs) were infected with the corresponding virus, followed by treatment with the candidate compounds. The levels of viral replication were then determined via quantitative real-time polymerase chain reaction (qRT-PCR) to evaluate the effects of each compound on the replication of its corresponding virus.

The cell-based *in vitro* results showed that a number of candidate compounds exhibited significant regulatory effects, either negatively or positively, on the replication or susceptibility of these representative viruses (Fig. 6a-e). Furthermore, several corresponding candidate compounds of HSV-1 and IAV were selected for further validation through *in vivo* animal experiments. As shown in Fig. 6f, the results suggests that GSK-461364 exhibited potent inhibitory activity against HSV-1 replication *in vivo*. Similarly, Nimesulide displayed favorable efficacy in suppressing IAV replication in mice (Fig. 6g).

**Figure 6.**
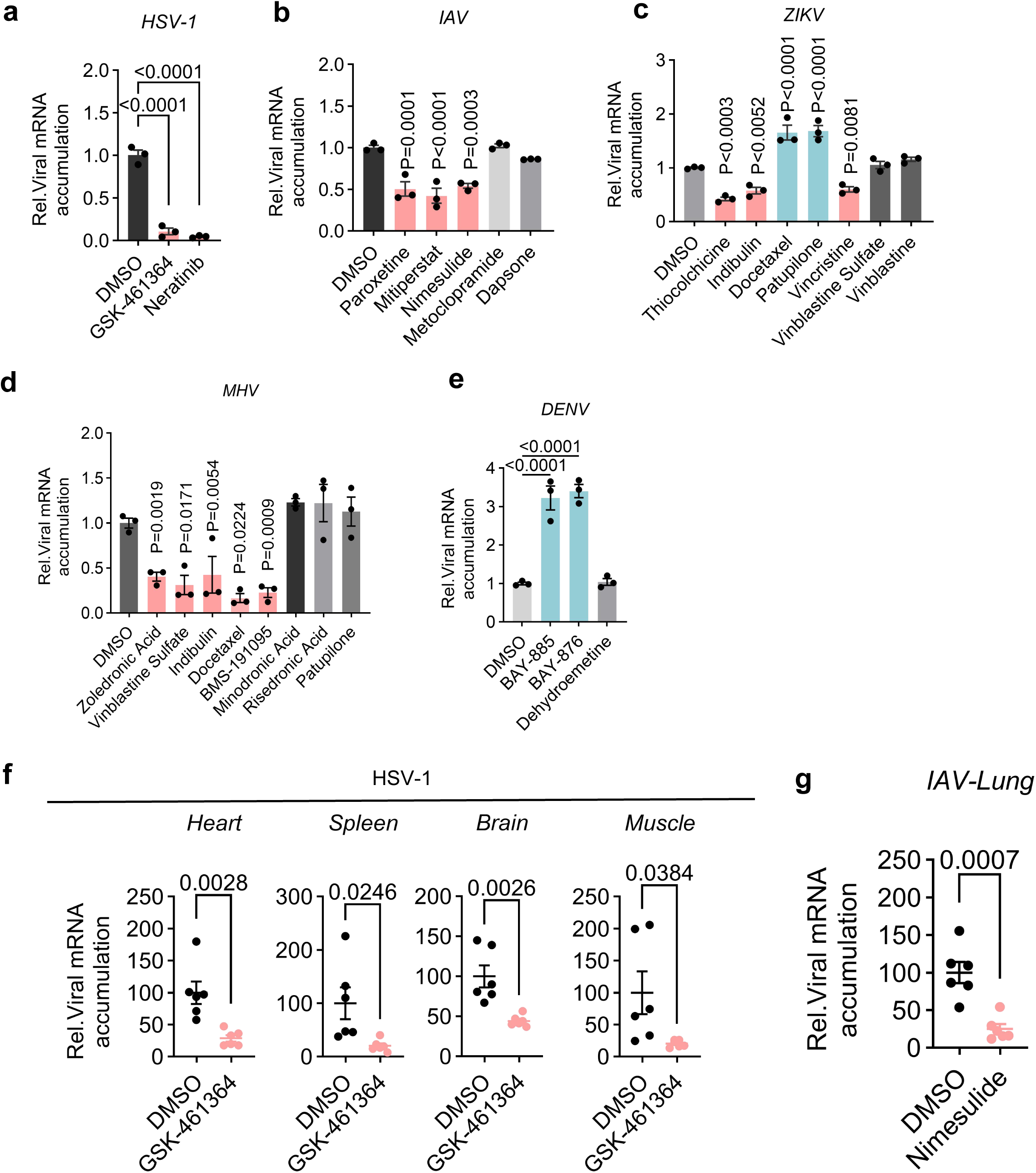
Compounds targeting selected VAHFs shows antiviral or proviral efficacies on the replication of multiple viruses in cells and mice. **a-e**, At 1 hr post infection (h.p.i) of indicated viruses, HeLa cells (a-c, e) or MLFs (d) were treated with 10 μM of the indicated candidate compounds. At 24 h.p.i., cells were collected and viral replication was measured by qRT-PCR with β-actin as the housekeeping control. Data were normalized to vehicle (dimethyl sulfoxide, DMSO) treated. Results for compounds that shows antiviral activity are displayed as pink bars, while those showing proviral activity are displayed as cyan bars. Those with no effect (i.e. no statistical significance) are represented by gray bars. Results for compounds exhibiting high cytotoxicity are not shown in the figure, and full details are available in the Supplementary Table. 5. **f,** Mice were treated with GSK-461364 or vehicle (DMSO) for 1 day prior to intravenous (i.v.) injection of HSV-1. At 72 d.p.i., total RNAs were extracted from heart, spleen, brain and muscle of infected mice, followedby qRT-PCR to measure viral RNA accumulation. **g,** Mice were treated with Nimeselide (10 mg/kg) or vehicle (DMSO) for 1 day prior to intranasal inoculation of IAV. Nimesulide or DMSO was administered once daily via i.p. injection for 2 consecutive days. At 48 d.p.i., total RNAs were extracted from lungs of infected mice, followed by qRT-PCR to measure viral RNA accumulation. Data were normalized to vehicle (DMSO) treated (f, g). Graphs show mean ± s.d.. Statistical significance is determined using one-way ANOVA (a-e) or student *t-*test (f, g).

All the tested compounds are categorized by the virus types, corresponding targets (VAHFs) and effectiveness, and are listed in Supplementary Table 5. Notably, since some VAHFs may not involve in viral susceptibility but mediate viral pathogenicity such as inflammation, cell death and tissue injury, pharmaceutical targeting these host factors may not affect viral replication in cells, but can still be authentic drug targets in treating viral diseases and associated symptoms. Therefore, our assay of measuring viral replication probably underestimated the capability of CellPulse in finding drug targets. Nevertheless, the performance of CellPulse in predicting drug targets and corresponding compound is impressive, which underscore the broad usability of CellPulse in drug discovery for diverse viral diseases.

### Experimental validation of VAHFs as therapeutic targets

After identifying a set of candidate compounds with promising therapeutic potentials based on the selected VAHFs, we aimed to further examine if these drug-corresponding host targets indeed exert regulatory effects on viral susceptibility.

For instance, NIMA-related kinase 2 (NEK2) is the corresponding host target for compounds GSK-461364 and Neratinib, which were confirmed to inhibit HSV-1 replication (Fig. 6a). Previous studies have reported that NEK2 is a serine/threonine protein kinase that plays a crucial role in cell cycle regulation, particularly during mitosis^23^. Of note, NEK2 has not been reported to involve in viral infection. To elucidate its role in HSV-1 replication, we constructed overexpressing plasmids and target-specific small interfering RNA (siRNA) for NEK2. HeLa cells and MLF cells were transfected with the aforementioned overexpressing plasmids or siRNAs, and qRT-PCR was then conducted to assess the impacts of their overexpression or knockdown on HSV-1 replication. As shown in Fig. 7a, compared with the empty vector-transfected control group, the overexpression of NEK2 significantly enhanced HSV-1 replication in HeLa and MLF cell lines. Conversely, silencing of NEK2 by specific siRNA remarkably suppressed HSV-1 replication relative to the nonsense siRNA-transfected control group. These observations were consistent with the antiviral phenotypes caused by its corresponding compounds, confirming the causal link between these host factors and HSV-1 replication regulation.

**Figure 7.**
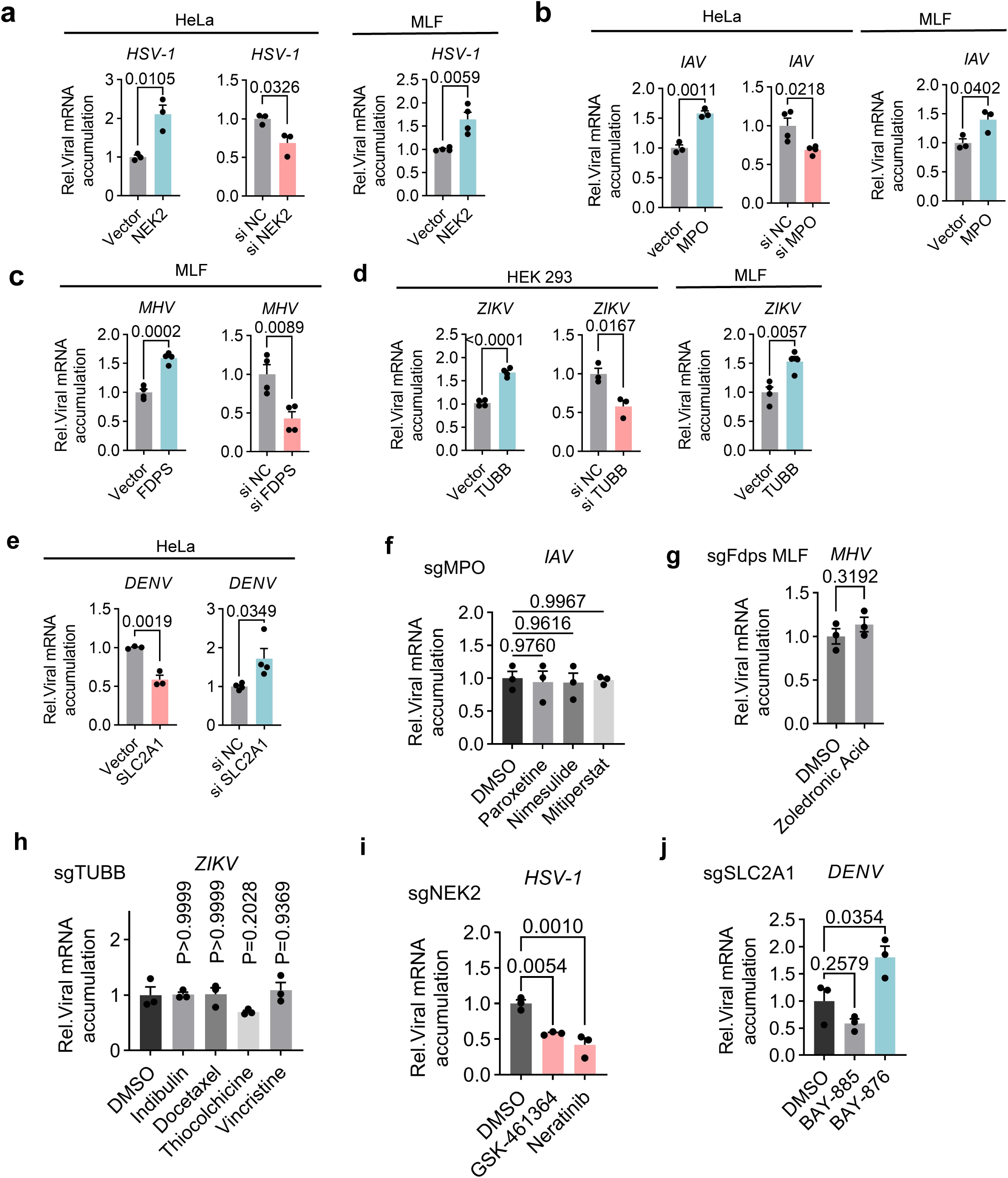
Experimental validation of selected VAHFs as therapeutic targets. **a-e,** Cultured HeLa cells, HEK293 cells, or MLFs were transfected with the plasmids expressing or siRNAs targeting the indicated genes. After 24 hours, cells were infected with indicated viruses. At 24 h.p.i., cells were collected and viral replication was determined via qRT-PCR with β*-*actin as the housekeeping control. Data were normalized to vector or siRNA negative control (siNC). **f-j,** At 1 h.p.i., HeLa cells with the knockout (KO) of indicated genes (f, h-j) or MLFs with the KO of *Fdps* (g) were treated with 10 μM of indicated compounds. At 24 h.p.i., cells were collected and viral replication was determined via qRT-PCR with β*-*actin as the housekeeping control. Data were normalized to vehicle (DMSO) treated sample. Graphs show mean ± s.d.. Statistical significance is determined using student *t-*test (a-e, g) or one-way ANOVA (f, h-j).

We further extended this validation strategy to other viruses, including IAV, MHV, DENV, and ZIKV. For each virus, overexpression plasmids and target-specific siRNAs were constructed for the host targets corresponding to drug compounds confirmed by us (Fig. 6a-e). And the consistency between the drug-induced phenotypes and the target-mediated effects on viral replication was systematically verified (Fig. 7a-e). As presented in Fig. 6b-d, several candidate compounds exhibited inhibitory activity against IAV, ZIKV, and MHV replication, respectively. Correspondingly, overexpression of their respective target factors, myeloperoxidase (MPO) for IAV, farnesyl diphosphate synthase (FDPS) for MHV, and tubulin beta (TUBB) for ZIKV, promoted viral replication, while knockdown of these targets exerted inhibitory effects on the replication of the corresponding viruses (Fig. 7b-d). Notably, drugs BAY-876 and BAY-885 were found to enhance DENV replication (Fig. 6e). Consistently, overexpression of their corresponding target, Solute Carrier Family 2 Member 1 (SLC2A1), suppressed DENV replication, whereas knockdown of these targets facilitated DENV replication (Fig. 7e), showing that SLC2A1 is a novel anti-DENV host factor. Since all these host genes had not been reported to link with viral infections, future studies should uncover the underlying mechanisms.

To validate that effective candidate compounds exert their pharmacological effects via the corresponding VAHFs, we designed specific single-guide RNAs (sgRNAs) for the key VAHFs, including NEK2, MPO, FDPS, TUBB, and SLC2A1, and constructed the corresponding gene knockout (KO) cell lines. We then performed viral replication assays in these KO cell lines. MPO-KO and FDPS-KO cells were infected with IAV and MHV, respectively, and treated with the candidate compounds targeting MPO (i.e. Paroxetine, Nimesulide, and Mitiperstat) or FDPS (Zoledronic acid). As shown in Fig. 7f and 7g, the inhibitory effects of these compounds on viral replication were almost eliminated in MPO- or FDPS-KO cells compared with that in wild-type (WT) cells (Fig. 6b and 6e). Moreover, TUBB-KO cells were infected with ZIKV and subsequently treated with TUBB-targeting compounds, including Indibulin, Docetaxel, Thiocolchicine and Vincristine. The results showed that all the examined compounds failed to suppress ZIKV replication in cells lacking the corresponding target (Fig. 7h and 6c). The loss of NEK2 dramatically attenuated the antiviral effect of GSK-461364 and Neratinib against HSV-1 (Fig. 7i and 6a), while the loss of SLC2A1 mostly abolished the ability of BAY-885 to enhance DENV replication, and partially attenuated the pro-viral effect of BAY-876 (Fig. 7j and 6d). These findings indicate that all these candidate compounds exert their pharmacological effects via the corresponding VAHFs.

Collectively, these results demonstrate a consistent causal relationship between drugs and their corresponding host targets in terms of their effects on viral replication, demonstrating that CellPulse is a powerful model in identifying novel host factors critical for viral life cycles and valuable targets for drug development.

## Discussion

This study presents VISTA, the first large-scale, AI-ready bulk RNA-seq perturbation atlas for viral infections, and CellPulse, a direction-aware transcriptomic foundation model trained on this dataset. Together, they establish a framework for systematically modeling host transcriptional responses, capturing mechanistic regulatory patterns, and enabling accurate virus type diagnostics and actionable insights into viral pathogenesis, host dependency factors, and potential therapeutic interventions.

Indeed, diagnosing virus types using real-world clinical samples poses significant challenges. First, many viral infections exhibit a transient nature, leading to extremely low viral nucleic acid concentrations in routinely collected specimens such as throat or nasal swabs, sputum, and blood, thereby imposes stringent requirements on sampling timing and frequently results in missed detection by conventional next-generation sequencing^24,25^. Second, the high mutation rate of viral genomes, coupled with the frequent fragmentation of viral nucleic acids compromise precise typing by sequence alignment algorithms^25–27^. Moreover, clinical samples commonly harbor nucleic acids from multiple co-infection or commensal pathogens, complicating the identification of true causative agent^25,27^, which typically requires laborious experimental validation based on Koch’s postulates^28^. Additionally, current nucleic acid-based detection/sequencing platforms are inherently limited in identifying or predicting novel or previously uncharacterized viruses^29^. In contrast, CellPulse circumvents these limitations by leveraging virus-induced perturbations in host gene expression to extract intrinsic, virus-specific transcriptional signatures (Virus Signatures). This approach enables accurate diagnosis of virus types solely on the basis of host response profiles and, by exploiting the causal relationship between Virus Signatures and associated pathologies, facilitating the discovery of key host factors and potential therapeutic targets.

Unlike existing medical AI models, which largely focus on imaging-based diagnosis^30^ or text-based clinical reasoning^31^, there is a critical gap in models that learn directly from host molecular responses^32^. While several single-cell transcriptomic models have demonstrated impressive predictive capabilities^7–12^, their reliance on scRNA-seq limits their applicability in real clinical scenarios due to sparsity, dropout bias, sample preparation constraints, and high cost, as well as stringent sample type requirement such as PBMCs and freshly acquired tissue biopsies. By contrast, bulk RNA-seq provides robust, cost-effective, quantitative measurements across diverse tissues and sample types, enabling models like CellPulse to capture subtle but biologically meaningful transcriptional changes. Our results demonstrate that transcriptional changes of genes with mid-to-low abundance, though subtle, carry meaningful signals that are critical for defining virus-specific regulatory signatures. The VISTA aggregates these responses at scale, enabling models to learn regulatory principles that are consistent across biological contexts. Indeed, before the VISTA, only the datasets based on scRNA-seq, which generates thousands of expression profiles in one sample, can include sufficient amount of data for training large models.

Despite the enormous complexity of gene regulation, which varies across tissues, stimuli, metabolic states, and individuals, gene expression changes in response to perturbation are not arbitrary. They reflect underlying biochemical and regulatory constraints that are based on fundamental biological principles and remain largely invariant across contexts. We posit that these constraints encode a perturbation grammar: a set of stable rules governing how genes co-vary when the system is challenged. Extracting such rules requires a modeling approach that does not memorize expression states, but instead learns how changes relate to other changes within the global regulatory network. CellPulse is designed not only to perform classification/diagnosis but also to learn gene co-regulation patterns under perturbation. The structured representation preserves both the magnitude and the direction of gene regulation changes. By explicitly modeling regulatory polarity, it avoids conflating activation with repression and enhances discriminatory power. The direction-aware attention mechanism emphasizes co-regulated gene modules and reveals mechanistic clusters consistent with known antiviral and virus-responsive pathways. Attention-based attribution identifies a large number of known and novel VAHFs, several of which we validated experimentally. These findings underscore the potential of the foundation model CellPulse not simply to diagnose diseases using conventional clinical samples such as whole bloods and throat or nasal swabs, but also to uncover mechanistic biological insights, thereby serving as hypothesis-generating engines for precision medicine and drug development.

Although this work focuses on viral infectious diseases, the principles underlying CellPulse are broadly applicable. Different diseases, such as autoimmune disorders, cancer, metabolic diseases, etc., exert distinct and reproducible regulatory pressures on gene networks. Because these regulatory patterns are governed by shared biochemical constraints, the same direction-aware modeling strategy could be extended to construct digital twins for diverse disease categories. Furthermore, the ability to systematically characterize transcriptional response patterns offers opportunities for drug repurposing, therapeutic optimization, and prediction of adverse effects. Given its high compatibility with diverse types of clinical samples, CellPulse represents a landmark toward developing digital-twin foundation models simulating infectious diseases and various other diseases for real clinical scenarios, which bridges high-dimensional omics data with diagnostics, therapeutics, and precision medicine.

## Methods

### Construction of VISTA dataset

The VISTA dataset was constructed through a unified, multi-stage processing pipeline designed to systematically integrate heterogeneous RNA-seq data from public repositories.

### Data collection

We systematically retrieved transcriptomic studies related to viral stimulation experiments from the NCBI Gene Expression Omnibus (GEO) database. To ensure biological relevance, we restricted data collection to human (*Homo sapiens*) and mouse (*Mus musculus*) samples. In total, VISTA incorporated 4,142 Gene Expression Series (GSEs), comprising 158,370 individual samples (GSMs) spanning a broad range of viruses, experimental designs, infection doses, and post-infection time-points.

### Data standardization

Public GEO datasets exhibit substantial heterogeneity in data formats, gene identifiers, and measurement scales, which poses major challenges for large-scale integration. To address these issues, we developed an automated standardization pipeline that converts all datasets into normalized, gene-aligned expression matrices with consistent Entrez gene identifiers and a unified species reference.

### Step A. Expression value normalization

Expression matrices in GEO may represent raw read counts, normalized abundance measures (e.g., Reads Per Kilobase of transcript per Million mapped reads [RPKM], Fragments Per Kilobase of transcript per Million mapped reads [FPKM], or Transcripts Per Million [TPM]), or log-transformed values. All entries were first converted to numeric format, with non-convertible values flagged as missing. For datasets lacking explicit normalization metadata, we employed heuristic checks (e.g., integer values for raw counts; non-integer values for normalized metrics). For non-normalized data, counts were converted to counts per million (CPM) using sample-specific total reads:

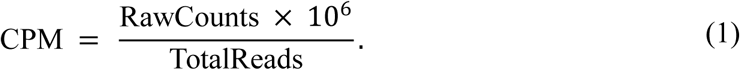

This procedure ensures comparability across samples while preserving relative expression differences.

### Step B. Probe-to-gene mapping

GEO platform annotation files were parsed to resolve probe-to-gene mappings across six commonly used identifier types: Entrez Gene IDs, GenBank accessions, RefSeq accessions, Ensembl IDs, UCSC symbols, and HGNC symbols^33–35^. For each annotation column, we evaluated the format-specific match rate, defined as the proportion of entries conforming to the expected syntax of a given identifier type. Columns with a match rate greater than 50% were retained, and the column with the highest match rate was selected as the primary gene identifier source. All non-Entrez identifiers were converted to Entrez gene IDs using official NCBI Gene mapping tables. For probes linked to multiple Entrez IDs (one-to-many relationships), the probe’s expression value was replicated across all associated genes. For multiple probes mapping to the same Entrez ID (many-to-one relationships), expression values from all probes were averaged.

### Step C. Human-mouse ortholog conversion

To enable robust integration of human and mouse transcriptomic data, we performed homology mapping between human and mouse Entrez IDs using gene symbols as the primary identifier. A total of 17,348 mouse genes were mapped to human orthologs. Species-specific genes lacking conserved counterparts (e.g., murine Glycam1) were systematically excluded to ensure functional comparability across species.

### Pairwise differential expression data generation

Rather than relying on predefined case-control labels, which are often missing or inconsistently defined across GEO submissions, we proposed an exhaustive pairwise comparison paradigm to generate large-scale differential expression (DE) data. Within each GSE, log2 fold changes were computed between all possible pairs of samples:

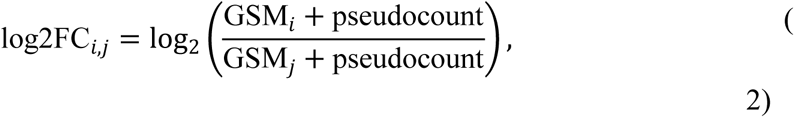

where GSM*_i_* and GSM*_j_* denote normalized expression vectors of sample *i* and *j* , respectively. A pseudo count (set to 0.01 in our calculations) is added to prevent zero values in the division and the logarithmic transformation. The pairwise comparison paradigm supports the integration of heterogeneous public datasets, even those lacking detailed metadata. Moreover, it enables large-scale transcriptional analysis without relying on predefined group labels, allowing for unbiased detection of regulatory changes across diverse experimental conditions, including dose responses, temporal progression, and combinatorial perturbations. Applying this procedure across all GSEs yielded 23,193,277 DE profiles covering 43,579 human genes.

### Viral perturbation data annotation

To enable systematic investigation of virus-specific host dependencies, we manually annotated biologically coherent experimental groups on 972 GSEs. Each group consists of biologically matched mock and viral-infection samples that are identical in experimental conditions except for viral exposure. Viral-infection samples were annotated with three key metadata fields: (1) virus taxonomy (e.g., IAV, SARS-CoV, Rhinovirus), (2) infection dose (quantified as multiplicity of infection, i.e., MOI, or plaque forming units, i.e., PFU), and (3) post-infection timepoints (e.g., 6 hours). Within each annotated group, log_2_ fold changes were computed between all control-treatment sample pairs, yielding 577,011 virus-associated DE profiles spanning 52 virus types.

### AI-ready data generation

To construct AI-ready data suitable for both self-supervised pretraining and task-specific fine-tuning, we applied two quality-control filters to every DE profile: For each DE profile, genes were first filtered based on absolute fold-change magnitude. Only genes satisfying a required threshold of log_2_FC ≥ were retained. This step removes weak or ambiguous responses and ensures that DE profiles emphasize biologically meaningful regulatory shifts. Second, each profile was required to contain at least two differentially expressed genes. This step guarantees that every training instance preserves a minimal degree of contextual structure, allowing the model to learn co-regulation patterns rather than degenerate single-gene fluctuations.

To examine how perturbation strength influences model performance and to provide datasets with different biological stringency, we constructed two variants of the corpus using commonly adopted fold-change thresholds: a permissive threshold of 1.2-fold ( ∣log _2_FC∣ ≥ 0.26 ) and a more stringent threshold of 1.5-fold ( ∣log _2_FC∣ ≥ 0.58 ). Applying the same filtering pipeline to both the full unlabeled pairwise-DE collection and the curated virus-perturbation subset yielded two harmonized datasets, denoted _1.2_ and _1.5_. These datasets were used consistently across both pretraining and fine-tuning stages, thereby enabling controlled comparisons of how the model behaves under varying levels of perturbation intensity while maintaining identical data construction principles throughout.

### CellPulse architecture

CellPulse introduces a direction-aware Transformer architecture for modeling gene co-regulation patterns from differential expression profiles. It represents perturbation-induced transcriptional responses as ranked gene sequences and integrates regulation direction as an inductive bias within the self-attention mechanism, enabling joint modeling of regulation magnitude, polarity, and coordinated transcriptional responses under biological perturbations such as viral infection.

### Base architecture

CellPulse consists of 12 sequential Transformer encoder layers. Each layer includes a multi-head self-attention module with 8 attention heads, operating over a hidden dimensionality of 512. Following each attention block, a two-layer feedforward network expands the representation to 2048 dimensions before projecting it back to 512, with GELU activation in between. Pre-layer normalization and residual connections are used to improve convergence and training stability.

### Input representation and embedding strategy

The inputs to CellPulse are constructed from DE profiles. For each DE profile, genes with an absolute log2FC value greater than *τ* are retained. These retained genes are then ranked by the absolute value of log2FC, emphasizing genes undergoing the most prominent changes. From this ranked list, a fixed number of top genes, determined by the model’s maximum input length, are selected to form the input sequence.

Each gene is represented by an integer index drawn from a predefined vocabulary of 43,579 human genes. Three additional special tokens are included: a classification token ([CLS]) is prepended to every sequence to provide a global sequence representation for downstream tasks, a masking token ([MASK]) used during pretraining, and a padding token ([PAD]) used to align sequence lengths within a batch. Given an input sequence 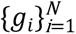 , where *N* denotes the total input length including the [CLS] and any [PAD] tokens, genes are embedded via a learnable embedding matrix *E_g_* ∈ ℝ^(43,579+3)×*d*^ . The three additional rows correspond to the three special tokens, and *d* = 512 denotes the hidden dimensionality. The gene identity embedding for *g_i_* is given by *E_g_*[*g_i_*]. To encode rank-order information derived from log_2_FC sorting, we further employ a learnable positional embedding matrix *E_p_* ∈ ℝ*^N^*^×*d*^. The final input embeddings are constructed as

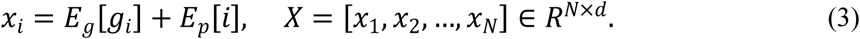

This formulation defines the core input representation of the CellPulse encoder, allowing the model to capture not only which genes are most altered, but also their relative ordering and contextual relationships within each DE profile.

### Direction-aware attention

Given that differential expression is inherently a relative measure and lacks absolute directionality and biological meaning (e.g., "up-regulation" is relative to a specific control) across arbitrary sample pairs, we introduced a directional co-regulation matrix *S* ∈ { − 1,0, + 1}*^N^*^×*N*^ . This symmetric matrix captures the directional concordance between each pair of genes in the input, defined as:

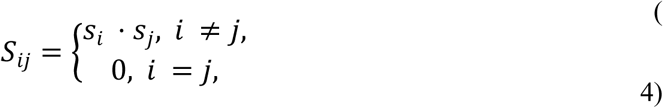

where *S_i_*, *S_J_* ∈ { − 1, + 1} denote the regulation direction (down- or up-regulated) for gene *i* and gene *j*, respectively. Thus, *S_ij_* =+ 1 implies concordant regulation (both genes up- or down-regulated), while *S_ij_* =− 1 indicates opposing regulatory directions. The diagonal entries are explicitly set to zero to prevent self-attention reinforcement through direction concordance, ensuring that the attention mechanism focuses on inter-gene regulatory relationships rather than trivial self-consistency. This directional co-regulation matrix *S* is incorporated directly into the scaled dot-product attention computation of each transformer layer:

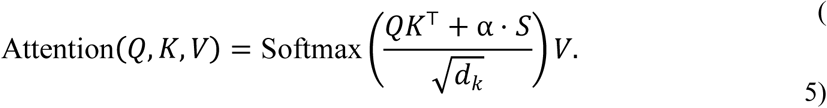

Here, 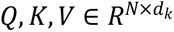 represent the usual query, key, and value matrices; *d_k_* is the dimensionality of the keys; and α is a learnable scalar parameter that modulates the influence of directional alignment. By incorporating *S* into the attention logits, the model is encouraged to preferentially attend to genes with concordant regulation, enhancing its capacity to learn biologically meaningful gene interactions.

The modified direction-aware attention mechanism is applied independently across all eight attention heads in each Transformer layer, with standard multi-head aggregation followed by position-wise feedforward networks. Residual connections and layer normalization are used throughout, consistent with Transformer best practices. This attention design enables CellPulse to attend not only to proximity in log2FC rank but also to the semantic alignment in regulatory directionality, providing a finer-grained representation of differential expression profiles.

### Optional direction-aware token augmentation

While the core CellPulse encoder models regulation direction through pairwise co-regulation in self-attention, the architecture also supports an optional token-level augmentation that explicitly encodes regulation polarity at the input level. In downstream tasks where a consistent reference condition is available and individual up- or down-regulation carries task-discriminative meaning, each gene token can be augmented with a discrete sign embedding representing its regulation direction. This augmentation does not alter the encoder structure and can be enabled selectively depending on the task and training objective. Its concrete instantiation and usage are described in the fine-tuning protocol.

### Pre-training protocol

We pre-trained CellPulse on the two AI-ready datasets, *V*_1.2_ and *V*_1.5_ , derived from the VISTA resource, aiming to learn robust representations of gene co-regulation under diverse biological perturbations. For each DE profile, genes were ranked by the magnitude of their absolute log2 fold change, and the top 2,048 genes were retained to conform to the model’s maximum input length.

Pre-training followed a contextual masking strategy analogous to masked language modeling. Specifically, 15% of gene tokens in each input sequence were randomly selected for masking, and the model was trained to reconstruct their identities based on the contextual co-regulation patterns of surrounding genes. Masked tokens followed the standard BERT masking procedure: 80% were replaced by a special [MASK] token, 10% were replaced by a random gene, and 10% remained unchanged. The training objective minimized the cross-entropy loss between predicted and true gene identities at masked positions. By forcing the model to infer missing genes from coordinated transcriptional contexts, this objective encouraged CellPulse to internalize stable co-regulation patterns among genes that tend to vary together across perturbations. Importantly, regulation direction was not encoded at the individual gene-token level during pre-training. Because DE profiles were constructed from arbitrary sample pairs without a fixed biological reference, gene-level up- or down-regulation was only meaningful in a relative sense. Accordingly, directional information was incorporated exclusively through the directional co-regulation matrix in the attention mechanism, allowing the model to learn regulation concordance patterns without introducing token-level sign embeddings.

Pre-training was performed using the Hugging Face Trainer framework with DeepSpeed optimization. The key hyperparameters were as follows: batch size per device, 20, learning rate, 2e-4, warm-up steps, 10,000, maximum sequence length, 2,048, and total training epochs, 10. The DeepSpeed configuration employed ZeRO optimization stage 2 with contiguous gradient storage and overlapping communication enabled, while optimizer offloading disabled.

### Fine-tuning on virus perturbation data

To adapt the pretrained CellPulse encoder for downstream analysis of viral perturbations, we fine-tuned the model on curated DE profiles derived from matched virus-infected and control samples. Each DE profile was labeled by viral category, enabling supervised learning of host transcriptional signatures associated with distinct viral perturbations.

Unlike the pre-training stage, fine-tuning was performed on DE profiles with a consistent biological reference (infected versus control), under which individual up-or down-regulation carries explicit biological meaning. Accordingly, CellPulse enabled token-level encoding of regulation direction during fine-tuning to further enhance task-specific discrimination. Specifically, each gene token was associated with a discrete sign indicator *S_i_* ∈ { − 1,0, + 1}, where +1 and −1 represent up- and down-regulation, respectively, and 0 corresponds to padding or the classification token. These sign values were mapped into categorical indices and embedded through a learnable matrix *E_s_* ∈ ℝ^3×*d*^. The fine-tuning input embeddings were constructed as

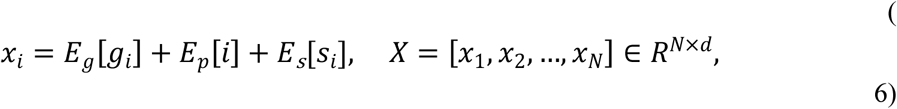

where *E_g_* and *E_p_* are identical to those used during pre-training. This additive integration allows CellPulse to condition contextual representations on regulatory direction without altering its pretrained co-expression structure.

The final sequence representation was obtained from the contextualized embedding of the [CLS] token, denoted ℎ_[CLS]_, which summarizes the transcriptional state of the input profile. Predictions were produced through a linear transformation followed by a softmax normalization:

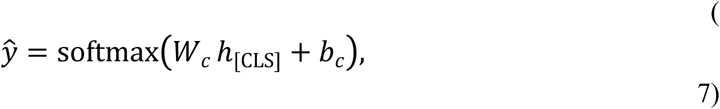

where *W_c_* ∈ ℝ*^d^*^×*c*^ and *C* is the number of output categories. Given the pronounced class imbalance inherent to viral perturbation classification, CellPulse was optimized using a class-weighted cross-entropy loss:

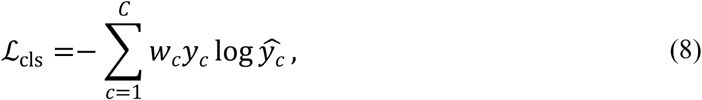

where *y_c_* denotes the one-hot encoded ground-truth label. The class weight *w_c_* is defined as

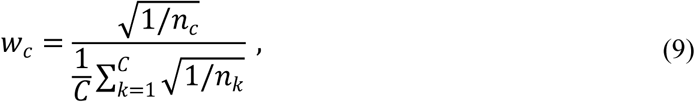

with *n_c_* representing the number of training samples in class *c* . This square-root inverse-frequency weighting mitigates dominance by majority classes while avoiding excessive amplification of rare classes, and the normalization ensures that the average class weight equals one.

Throughout fine-tuning, the direction-aware attention mechanism remained active, with the directional co-regulation matrix integrated into each self-attention computation. By jointly leveraging pairwise co-regulation biases and token-level regulation polarity, CellPulse learned task-specific decision boundaries while preserving biologically meaningful gene-gene interaction patterns underlying host-virus responses.

Fine-tuning was conducted using the Hugging Face Trainer framework with class-weighted loss. The first two encoder layers were frozen to preserve pretrained representations while allowing task-specific refinement. Hyperparameter optimization was performed using Ray Tune with a HyperOpt search strategy, exploring learning rate, weight decay, scheduler type, and warm-up steps. The final model was selected according to validation macro-F1 performance.

### Attention-based attribution

To evaluate whether CellPulse captures biologically meaningful regulatory structure and to identify host factors critical for virus-induced perturbations, we conducted a large-scale attention-based attribution analysis across viral classes. In CellPulse, the [CLS] token aggregates the global regulatory representation of each DE profile; therefore, the self-attention weights linking the [CLS] token to individual gene tokens provide a model-native measure of each gene’s relative contribution to the inferred perturbation pattern. Intuitively, higher [CLS] to gene attention indicates a stronger role for that gene in shaping the model’s interpretation of the regulatory contrast.

For each viral class, we randomly sampled up to 10,000 DE profiles (using all available profiles for classes with fewer instances), yielding a total of 76,038 profiles. Each profile was passed through the trained model, and the final-layer self-attention weights were extracted. To obtain a stable attribution signal, attention was averaged across all heads. Formally, let *A*^(*L*)^ ∈ ℝ*^H^*^×*T*×*T*^ denote the last-layer attention tensor, where *H* is the number of heads and *T* is the sequence length. The head-averaged attention matrix is defined as

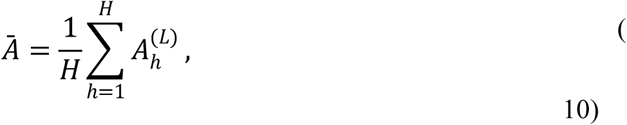

Let *g_i_* be the *i*-th gene token in the profile *j* . The profile-level gene importance score is then

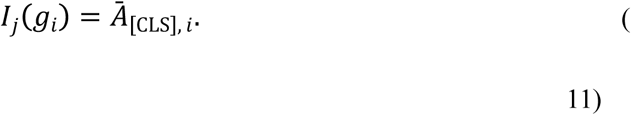

This quantity represents how strongly the model attends to *g_i_* when constructing its global regulatory representation. The resulting gene importance scores were then aggregated at the viral-class level. For a given viral class *c*, gene importance scores were averaged across all DE profiles belonging to that class. Let *n_c_* be the number of profiles associated with class *c*. The class-level importance score for gene *g_i_* is

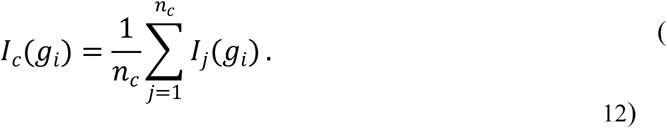

These class-level scores were used to rank genes by their recurrent importance within each viral category. These high-ranking genes represent candidate host factors whose regulatory perturbation patterns are most informative for distinguishing that viral infection.

This attribution framework leverages CellPulse’s internal regulatory weighting mechanism to reveal host genes that the model systematically relies upon when interpreting viral perturbations, enabling a biologically grounded interpretation of virus-host regulatory dependencies.

### Gene Ontology analysis (GO analysis)

GO analysis terms biological process was performed using R language and the *clusterProfiler* package^36–39^, with the main parameter as follows: OrgDb: org.Hs.eg.db; pvalueCutoff: 0.05; pAdjustMethod: fdr; ont: BP. The top 20 GO terms were selected for visualization, which was implemented using the *ggplot2* package^40^.

### Cell culture

HeLa, HEK293, Vero E6, C636, MDCK and L2 cells were commercially obtained from the American Type Culture Collection (ATCC). Mouse lung fibroblasts (MLFs) were isolated from mouse lungs and subsequently immortalized. Briefly, lungs were minced and digested in calcium- and magnesium- containing Hanks’ balanced salt solution (HBSS) buffer (Yeasen) supplemented with type II collagenase (5 mg/mL; Yeasen) and DNase I (20 μg/mL; Thermo Fisher) for 4 hours at 37 ℃. Cell suspensions were filtered through progressively smaller cell strainers (100-40μm) and filtered cells were plated with DMEM containing 10% FBS, 2mM L-glutamine, 100 U/mL penicillin, and 100 μg/mL streptomycin. After 1 hour incubation, adherent MLFs were rinsed with HBSS and cultured for the subsequent experiments. The immortalized MLFs were performed by using commercially immortalized kit (LINGSIBIO, LJSCI-002), following the manufacturer’s instructions. The VAHFs-KO-HeLa/MLFs were constructed via CRISPR/Cas9 system. Without otherwise specified, cells were cultured in DMEM (Gibco) containing 10% FBS (Gibco) and 1% streptomycin-penicillin (Gibco) at 37 ℃ in a 5% CO2 incubator.

### Viruses

MHV (Strain A59) were amplified in L2 cells. IAV (A/Puerto Rico/8/1934, H1N1) were amplified in chick embryo and MDCK cells. HSV-1 (Strain F) were propagated at a low MOI in Vero E6 cells. ZIKV and DENV were amplified in C6/36 cells, which were cultured in RPMI 1640 (Gibco) containing 10% FBS (Gibco) and 1% streptomycin-penicillin (Gibco) at 27 ℃ in the absence of CO2.

Titers were determined in the corresponding cells by plaque assay. Before infection with viruses, the medium was changed to DMEM containing 2% FBS without any antibiotic. Virus-related experiments were conducted in Biosafety Level-2 (BSL-2) labs at Wuhan Institute of Virology, CAS.

### Plasmids and siRNAs

Plasmids construction was described previously^41^. Briefly, the DNA fragment of VAHFs were synthesized by Hecegene Technology Co., Ltd (Wuhan, China) and ligated to the pRK5-FLAG vector by homologous recombination enzyme (ABclonal, RK21020) following the manufacture’s instruction. All plasmids were confirmed by DNA sanger sequencing.

The small-interfering RNAs (siRNAs) were synthesized by GeneCreat (Wuhan, China) and were transfected with Lipofectamine 2000 reagent into cells cultured as 50-70% confluence. All the siRNA sequences used here are listed in Supplementary Table 6.

### Plaque assays

Plaque assays were performed on corresponding cells in a 12-well plate infected with a 10-fold serial dilution of viruses. After 1 hour incubation for absorption, the supernatants were removed and infectious cells were washed with pre-warmed PBS twice. Subsequently, cells were overlaid with 1% low-melting-point agarose in DMEM containing 2% FBS. After further incubation for 48 hours, cells were fixed with 4% formaldehyde and stained with 0.5% crystal violet to visualize and count the plaques.

### CRISPR/Cas9-based gene knockout

The sgRNAs were used to knockout the corresponding VAHFs in HeLa cells and MLFs as previously described^42^. Briefly, synthesized sgRNA oligonucleotides were cloned into the pLentiCRISPR-V2 vector using T4 DNA ligase (Thermo Fisher). All the pLentiCRISPR-sgRNA plasmids were confirmed by DNA sanger sequencing. The pLentiCRISPR-sgRNA plasmids were co-transfected into HEK 293 cells together with the lentivirus packaging plasmids psPAX2 and pMD2.G to produce lentiviral particles. These lentiviruses were subsequently used to stably transduce HeLa cells or MLFs. HeLa cells and MLFs were cultured for 3 days and selected by puromycin (2 μg/mL for HeLa cells and 1.5 μg/mL for MLFs, InvivoGen). To generate clonal KO lines, single cell cloning was seeded on to the 96-well plates. The resulting clones were confirmed by western blotting and DNA sanger sequencing. All sgRNA sequences used here are listed in Supplementary Table 6.

### Quantitative real-time PCR (qRT-PCR)

Cells were harvested and total RNA were isolated using a cell total RNA isolation kit (Foregene, RE-03111). The first-strand cDNA was reverse-transcribed with 1st strand cDNA Synthesis (Takara, D6110A). Gene expression was examined with a Bio-Rad SFX connect system by a qPCR SYBR Green Master Mix (Yeasen, 11198ES03). Data were normalized to the expression of the gene encoding β-Actin. All the qPCR primers used here are listed in Supplementary Table 7. The assays were conducted using ABI QuantStudio Q3 real-time PCR system and the data were collected using QuantStudio design and analysis software v1.4.

### Compounds screening and target VAHF(s) validation assays

The candidate compounds were dissolved in dimethyl sulfoxide (DMSO) to a final concentration of 10 mM. HeLa cells seeded in 48-well plates were infected with 1 MOI of multiple virus (HSV-1, IAV, DENV, ZIKV) and MLFs seeded in 48-well plates were infected with 1 MOI of MHV. At 1 hour post infection (h.p.i.), the supernatants were removed, and the infected cells were washed with pre-warmed PBS twice followed by incubation with compounds diluted in fresh DMEM supplemented with 2% FBS to a final concentration of 10 μM. At 24 h.p.i., cells were harvested and viral replication levels were measured via qRT-PCR. All candidate compounds were purchased from TargetMol (Shanghai, China).

For the transfection of the target VAHFs, corresponding cells seeded in 24-well plates were transfected with either 100 nM small interfering RNA (siRNA) or 800 ng plasmids. The transfection complexes were prepared by mixing the nucleic acid constructs with 2.5 μL of Lipofectamine 2000 reagent and 100 μL of Opti-MEM reduced-serum medium (Gibco, 22600134) prior to addition to the cell cultures. At 24 hours post-transfection (h.p.t.), cells were inoculated with the corresponding viruses at a MOI of 0.1. At 24 h.p.i., cells were harvested and viral replication levels were measured via qRT-PCR.

### Mouse experiments

Six-week-old C57BL/6J mice were intraperitoneally (i.p.) injected with Nimesulide (10 mg/kg) or DMSO vehicle. After 24 hours, mice were challenged with IAV (1 x 10^4^ PFU/ per mouse) via intranasal inoculation. Nimesulide or DMSO vehicle was administered once daily via i.p. injection for 2 consecutive days. Mice were euthanized at 2 days post infection (d.p.i) and lung samples were harvested for subsequent analysis.

Six-week-old C57BL/6J mice received a single dose of GSK-461364 (20 mg/kg) or DMSO vehicle via intraperitoneal (i.p.) injection. After 24 hours, mice were challenged with HSV-1 (1 x 10^7^ PFU/ per mouse) via intravenously (i.v.) injection. Mice were euthanized at 3 d.p.i., and heart, spleen, brain and muscle samples were harvested for subsequent analysis.

All mice were purchased from Beijing Vital River Laboratory Animal Technology Co., Ltd. and housed in the animal facility at the Center for Experimental Animals of Wuhan Institute of Virology, CAS. The mice were maintained specific-pathogen free (SPF) conditions on a 12/12-hour light/dark cycle with sterile standard rodent chow and water ad libitum. All animal experiments were conducted in strict accordance with the guidelines of Regulation on the Administration of Laboratory Animals (Ministry of Science and Technology) and were approved by the Institutional Animal Care and Use Committee (IACUC) of Wuhan Institute of Virology, CAS.

### Quantification and statistical analysis

GraphPad Prism (v10.6.1) was used for the statistical analyses. The sample number of repeats are indicated in the respective figure legends. Statistical significance was calculated by one-way analysis of variance (one-way ANOVA) or student *t-*test analysis, as indicated in the corresponding figure legends.

### Data availability

This study did not generate new unique reagents. All data are available in the main text or supplementary materials. All bulk RNA-seq data integrated into VISTA were obtained from publicly accessible GEO database (https://www.ncbi.nlm.nih.gov/geo/), and corresponding GSE accession numbers are list in Supplementary Table 8. The VISTA dataset has been deposited to the Open Archive for Miscellaneous Data (OMIX) in National Genomics Data Center (NGDC) under a secure private link (https://ngdc.cncb.ac.cn/omix/preview/62avCncb) for peer review. The VISTA dataset will be made publicly available under accession number OMIX ID: OMIX015013 and BioProject ID: PRJCA058035 upon manuscript acceptance.

### Code availability

The source code developed for this study, including data pre-processing pipelines for VISTA construction, CellPulse model implementation, pre-training, fine-tuning and evaluating scripts, attention-based attribution analysis, and in silico drug screening workflows, has been deposited in a restricted-access Zenodo repository. Pretrained and fine-tuned model weights are included. The training pipeline was implemented using the Hugging Face Trainer framework, with certain implementation utilities developed with reference to open source Geneformer codebase. All model architectures, biological tasks, and experimental designs are original to this work. During peer review, the repository is accessible to editors and reviewers via a secure private link:

https://zenodo.org/records/18616493?preview=1&token=eyJhbGciOiJIUzUxMiJ9.eyJpZCI6IjM1OWEwMjBmLTNmMTktNDM0ZS05MjM0LTY1ZWU4NWE2YWY3MCIsImRhdGEiOnt9LCJyYW5kb20iOiI1Y2U2YjkwNjc4OTAyMWNiNzYyMzVmNGJkNTkxZDQ5YyJ9.KUBgELK9zOI9iwV0d3whGp1hlH6WfUK2chr4kbFSYrA0OYZxu8jzMyNBy7lFmsL2dPghVsz14-O_e4IdR6W_uw. The repository will be made publicly available under the MIT license upon manuscript acceptance.

Further information and requests for resources and reagents should be directed to and will be fulfilled by Xi Zhou.

## Acknowledgments

We thank all the members of Zhou group, Wu group, and the WIV-IS Innovation Center of Artificial Intelligence for Disease and Therapy Research for their support. This work was supported by the National Natural Science Foundation of China (82550129 to X.Z., 8252200663 and 82472274 to Y.R.).

## Author contributions

X.Z., Y.W., Y.R. and L.Z. designed and supervised the studies. D.L. and X-X.Z. led the development and construction of the VISTA database with the help of D.X.. D.L. and L.Z. led the overall model development. X-X.Z. performed most of the validation experiments with the help of D.X., J.L. and X.X. Y.R., X-X.Z., and D.X. analyzed the data. X.Z., Y.W., Y.R., L.Z., D.L. and X-X.Z. wrote the manuscript with input from all the authors.

---

**Supplementary Figure S1.**
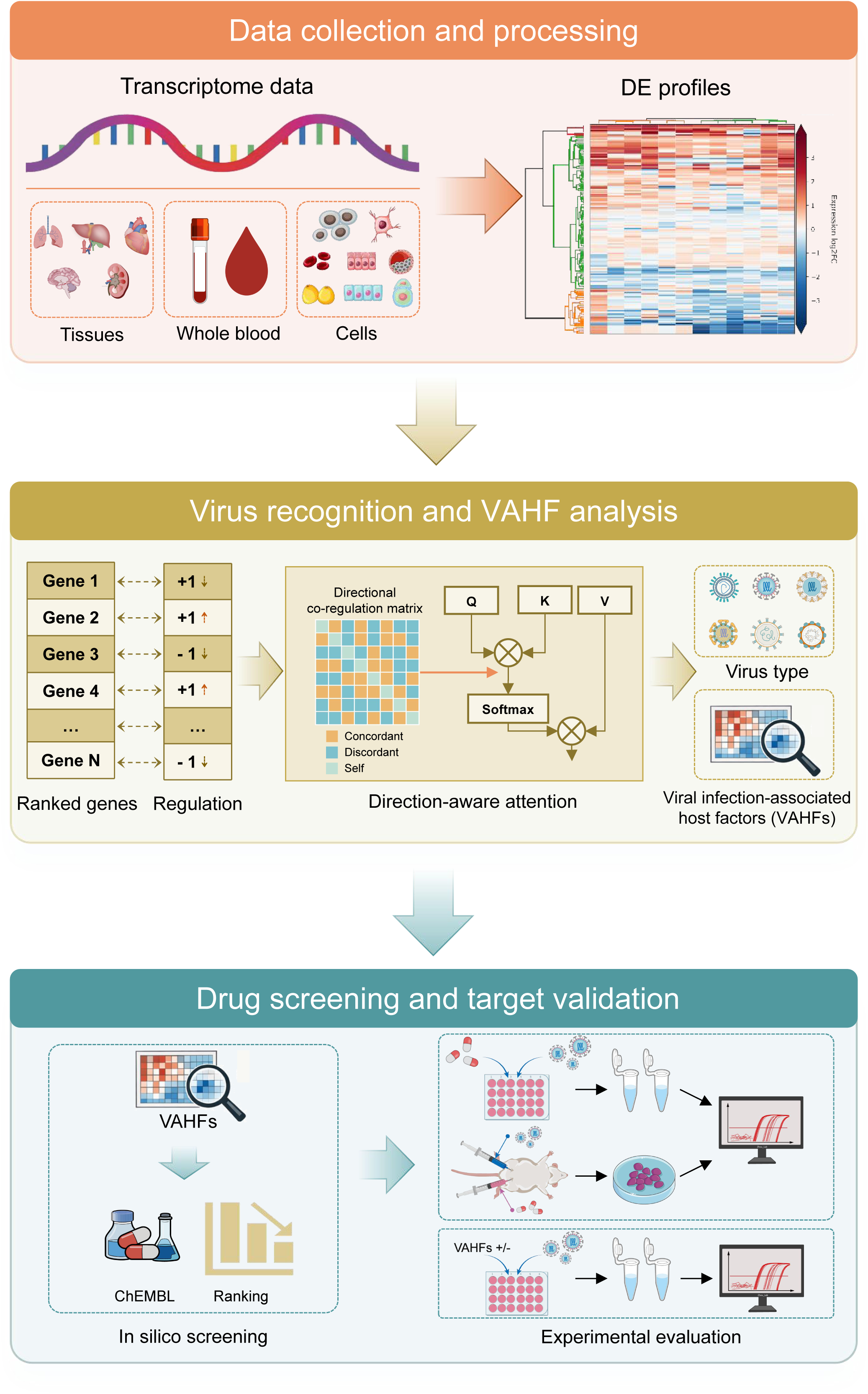
Schematic workflow of the study. (1) Data collection and processing to build VISTA: Bulk RNA-seq transcriptome data from diverse clinical specimens and laboratory samples (e.g., tissues, whole blood, cells) were curated and processed to generate differential expression (DE) profiles, forming the VISTA. (2) Virus recognition and VAHF analysis: CellPulse ingests DE profiles as ranked gene lists with direction vectors (+1 for upregulation, -1 for downregulation). A directional co-regulation matrix is integrated into the self-attention mechanism to prioritize concordant regulatory patterns, enabling virus type identification and inference of VAHFs via attention-based attribution. (3) Drug screening and target validation: Top VAHFs are mapped to the ChEMBL database for *in silico* compound ranking. Candidate compounds undergo *in vitro* (cell-based) and *in vivo* (mouse models) validation, with VAHF overexpression and loss-of-function (siRNA knockdown, CRISPR/Cas9 knockout) experiments confirming the causal roles of VAHFs in viral susceptibility.

**Supplementary Figure S2.**
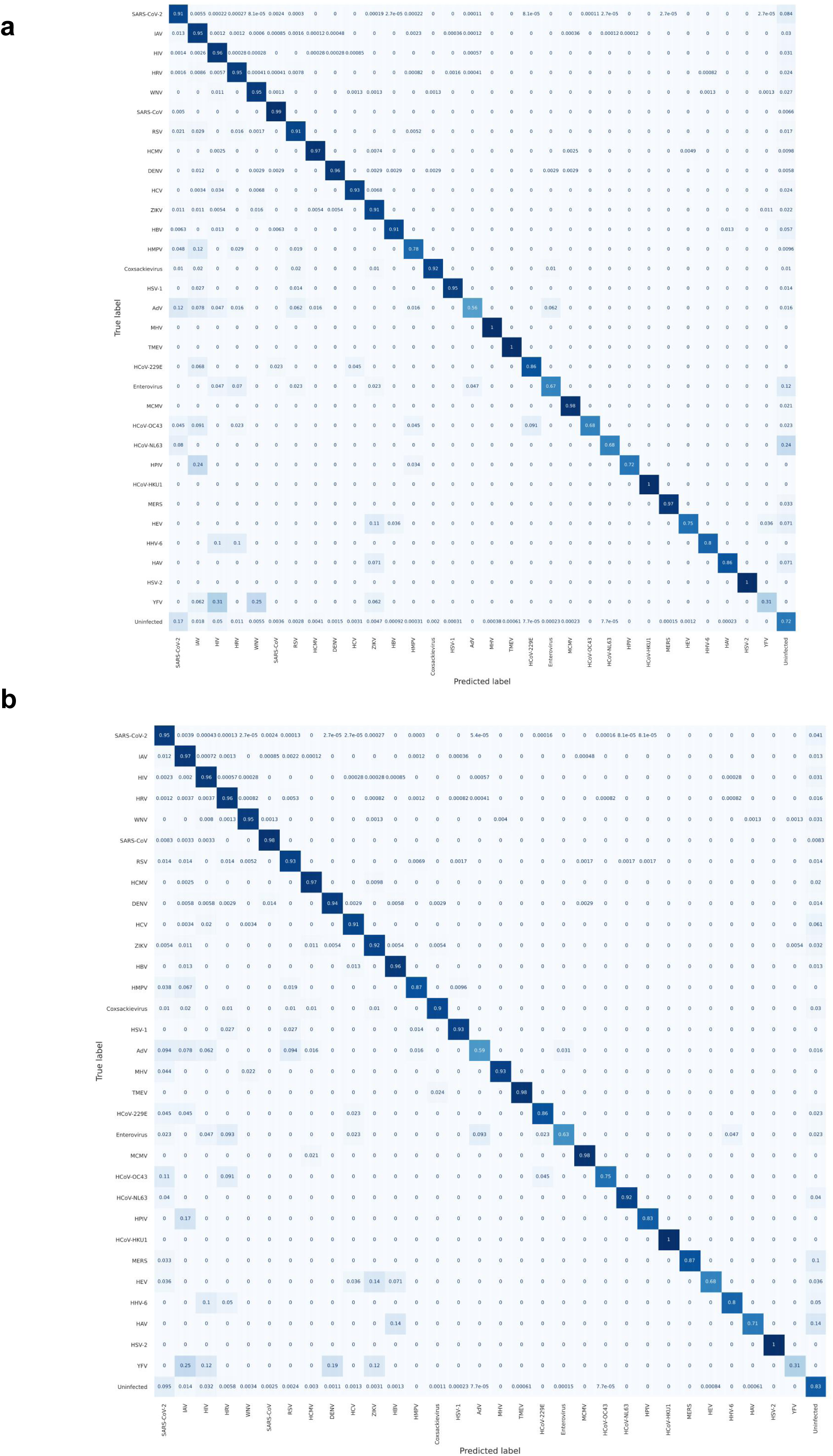
Performance of BERT and CellPulse on the 1.5-fold dataset V_(1.5). **a**, Normalized confusion matrix for the BERT baseline. **b**, Normalized confusion matrix for CellPulse.

**Supplementary Figure S3.**
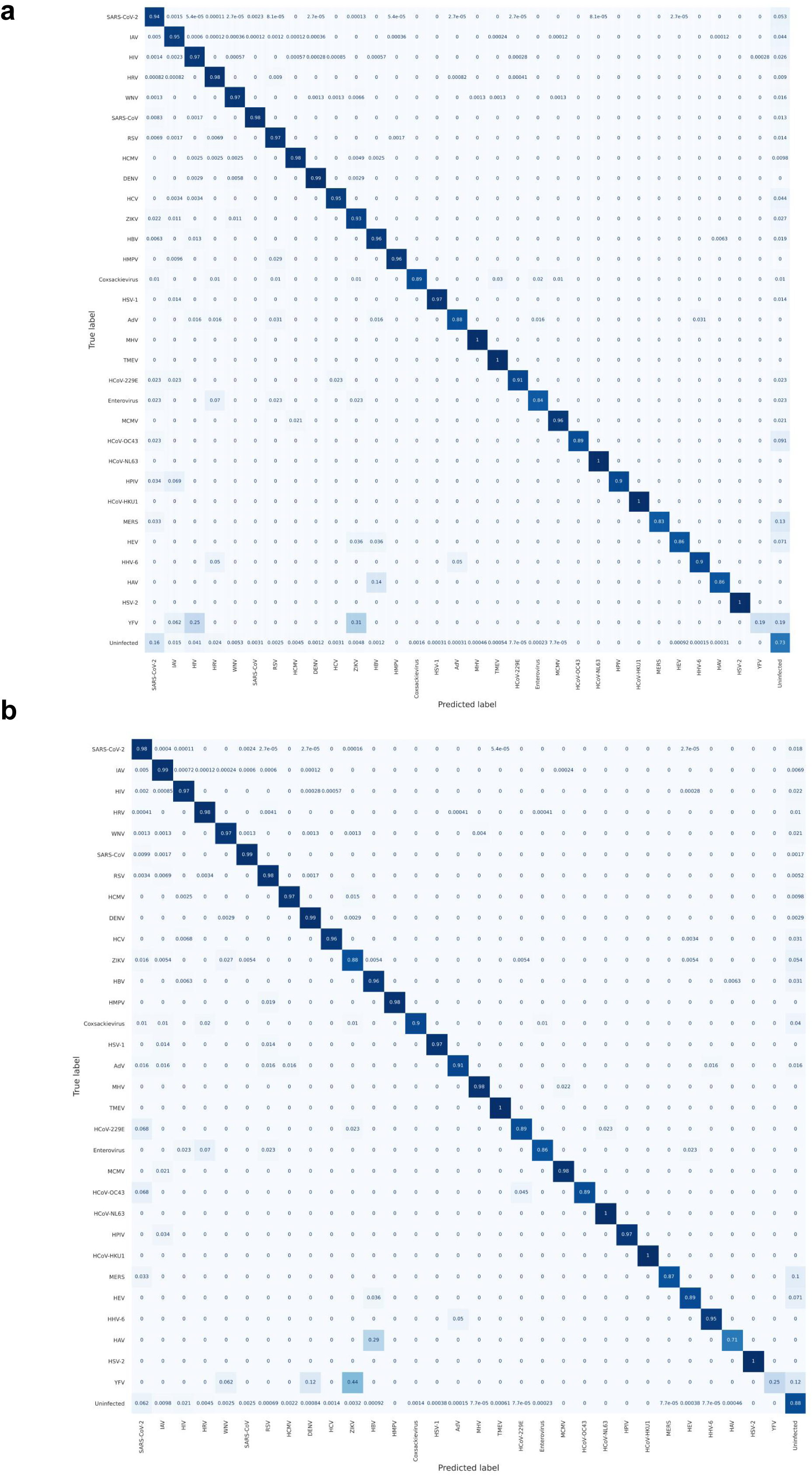
Performance of BERT and CellPulse on the 1.2-fold dataset V_(1.2). **a**, Normalized confusion matrix for the BERT baseline. **b,** Normalized confusion matrix for CellPulse.

**Supplementary Figure S4.**
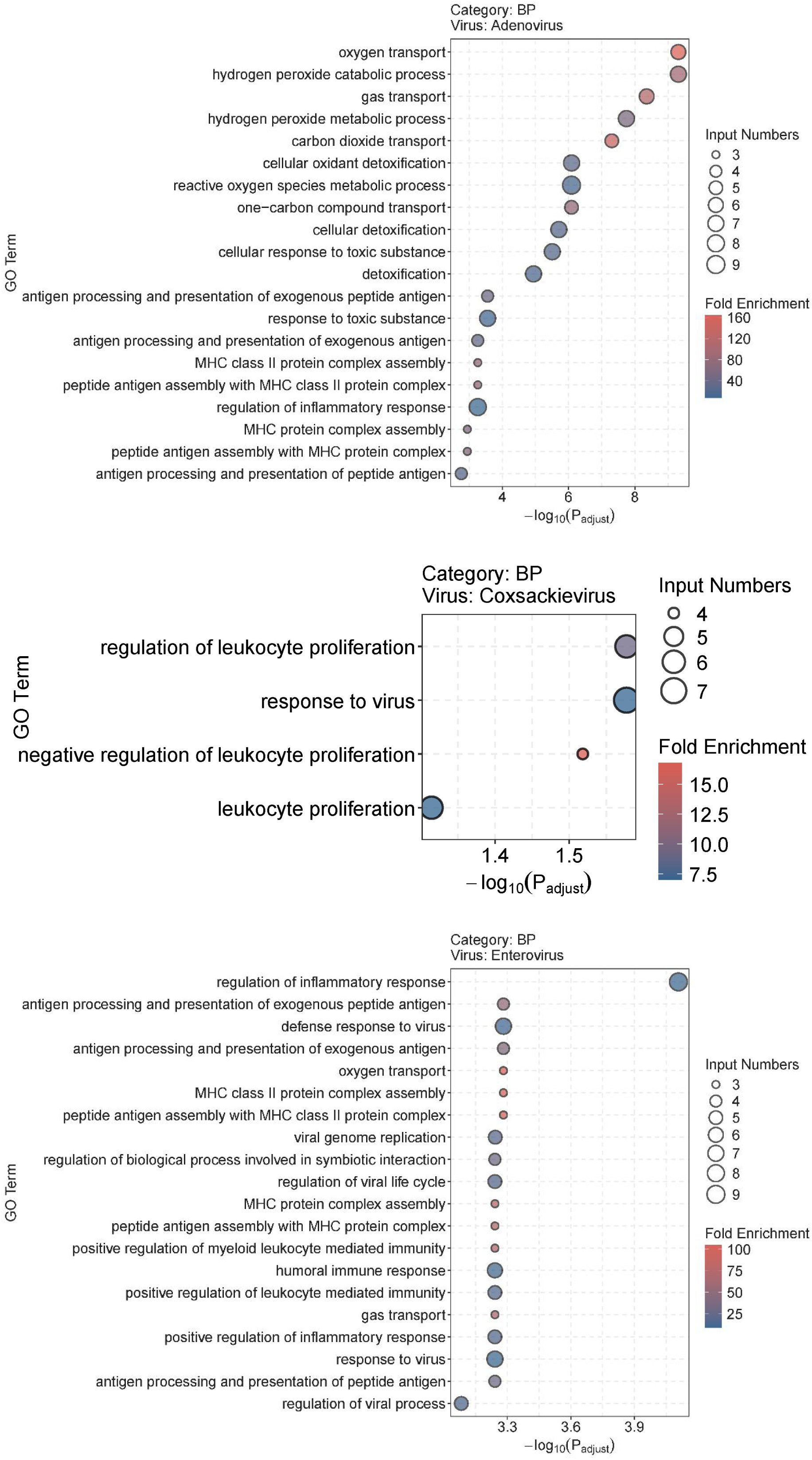
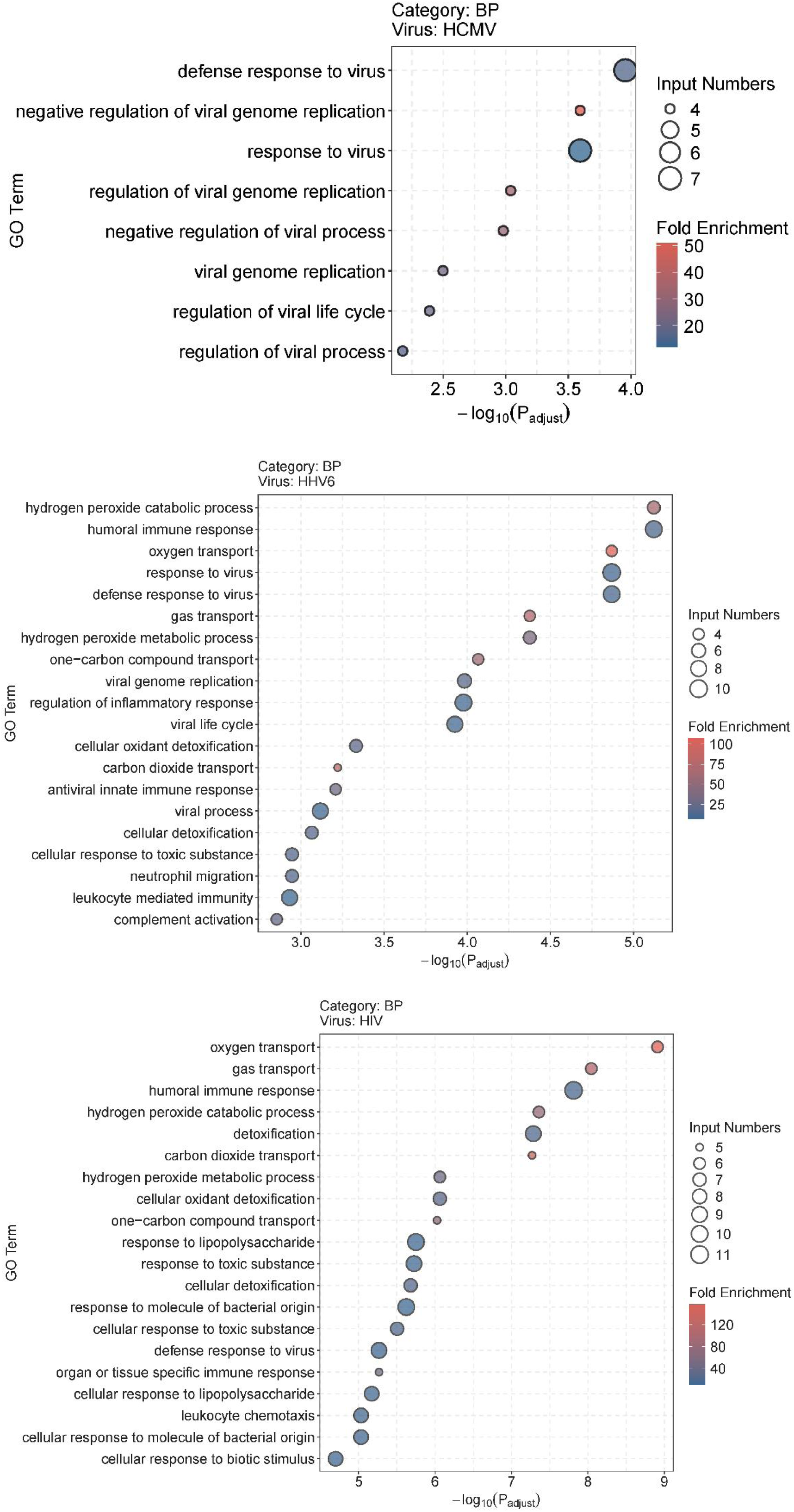
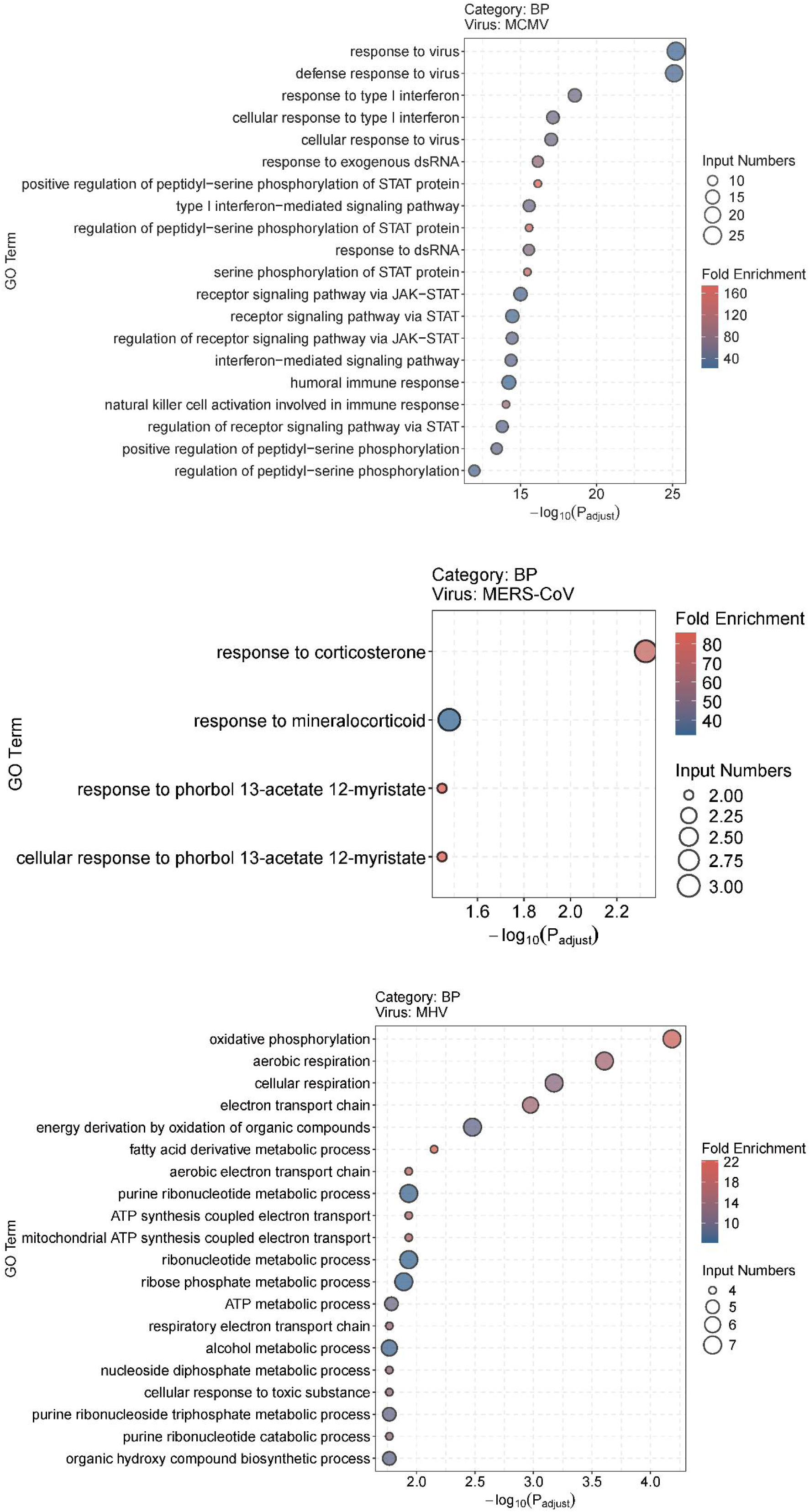
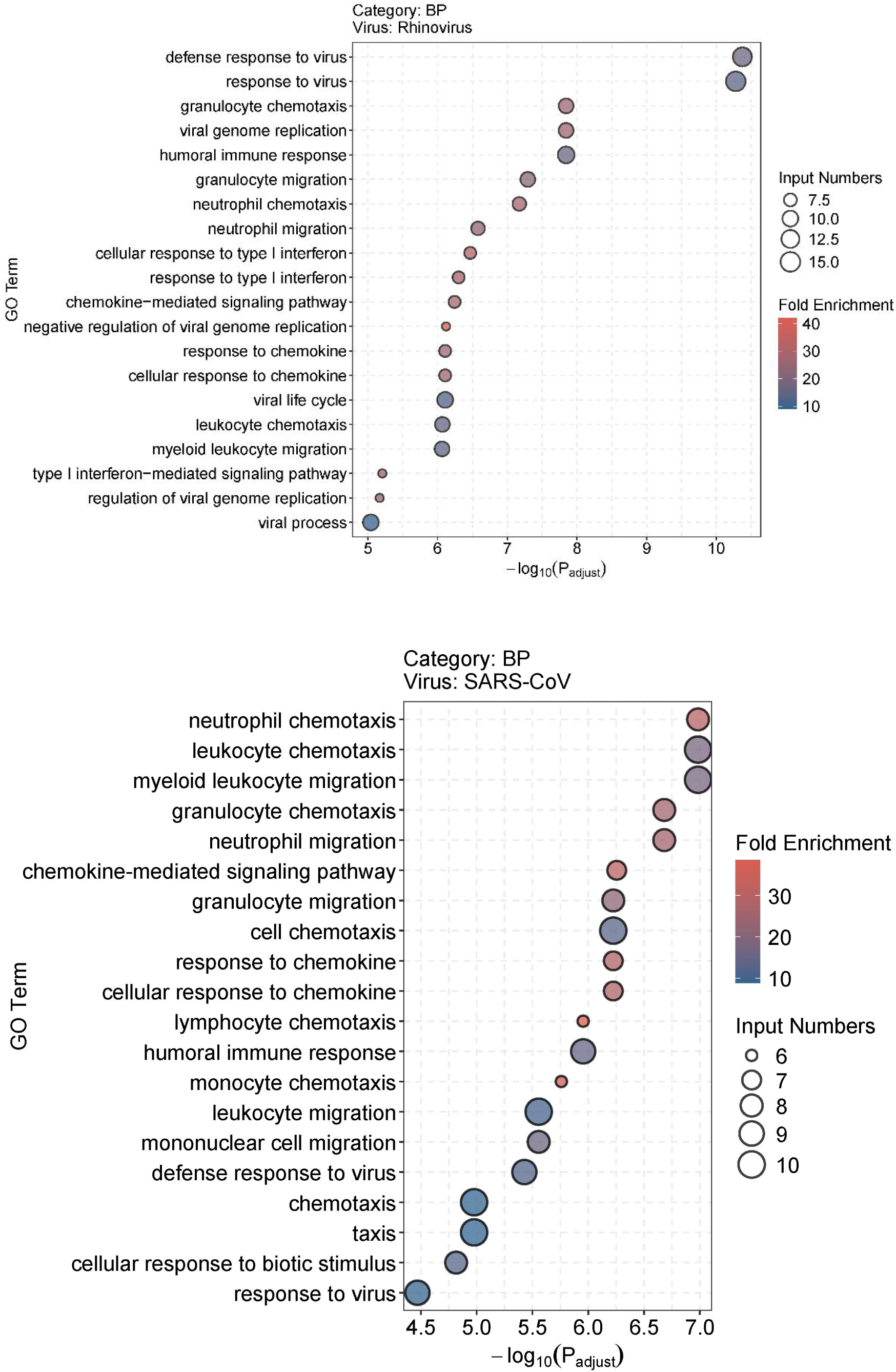
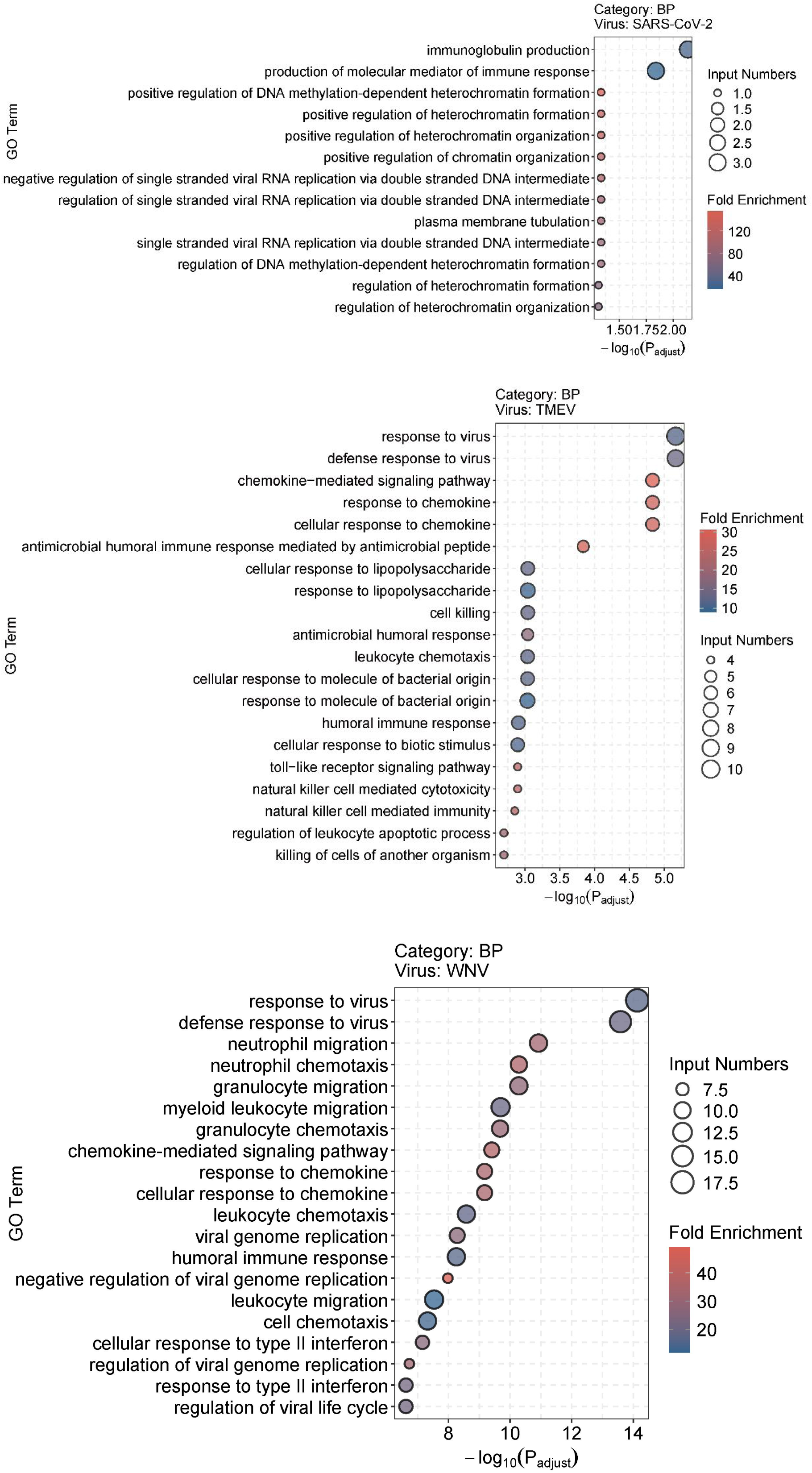
Bubble plot of the top significantly enriched GO on BP terms associated with the gene set corresponding to the top 50 VAHFs of individual virus. GO enrichment analysis was conducted using an FDR threshold of < 0.05 for statistical significance. Bubble size represents the number of genes annotated to each term, and color intensity indicates the fold enrichment.

